# The 14q32.31 *DLK1-DIO3 MIR300 tumor suppressor* promotes leukemogenesis by inducing cancer stem cell quiescence and inhibiting NK cell anti-cancer immunity

**DOI:** 10.1101/680108

**Authors:** Giovannino Silvestri, Rossana Trotta, Lorenzo Stramucci, Justin J. Ellis, Jason G. Harb, Paolo Neviani, Shuzhen Wang, Ann-Kathrin Eisfeld, Christopher Walker, Bin Zhang, Klara Srutova, Carlo Gambacorti-Passerini, Gabriel Pineda, Catriona H. M. Jamieson, Fabio Stagno, Paolo Vigneri, Georgios Nteliopoulos, Philippa May, Alistair Reid, Ramiro Garzon, Denis C. Roy, Moutua-Mohamed Moutuou, Martin Guimond, Peter Hokland, Michael Deininger, Garrett Fitzgerald, Christopher Harman, Francesco Dazzi, Dragana Milojkovic, Jane F. Apperley, Guido Marcucci, Janfei Qi, Katerina Machova-Polakova, Ying Zou, Xiaoxuan Fan, Maria R. Baer, Bruno Calabretta, Danilo Perrotti

## Abstract

Drug-resistance of tumor-initiating cells, impaired NK cell immune-response, PP2A loss-of-function and aberrant miRNA expression are cancer features resulting from microenvironmental- and tumor-specific signals. Here we report that genomic-imprinted *MIR300* is a cell context-independent dual function tumor suppressor which is upregulated in quiescent leukemic stem (LSC) and NK cells by microenvironmental signals to induce quiescence and impair immune-response, respectively, but inhibited in CML and AML proliferating blasts to prevent PP2A-induced apoptosis. *MIR300* anti-proliferative and PP2A-activating functions are differentially activated through dose-dependent CCND2/CDK6 and SET inhibition, respectively. LSCs escape PP2A-mediated apoptosis through TUG1 lncRNA that uncouples and limits *MIR300* functions to cytostasis by regulating unbound-*MIR300* levels. Halting *MIR300* homeostasis restores NK cell activity and suppresses leukemic but not normal hematopoiesis by eradicating nearly all LSCs. Thus, *MIR300* tumor suppressor activity is essential and therapeutically important for LSC-driven leukemias.

## INTRODUCTION

Functional loss of protein phosphatase 2A (PP2A) tumor suppressor activity is a therapeutically-relevant cancer feature which is essential for cancer stem cells (CSCs) maintenance, tumor growth/progression and activation of natural killer (NK) cell proliferation and cytotoxic activity^1–3^. Microenvironment and tumor-specific factors (e.g. cytokines, oncoproteins) inhibit PP2A activity through the activation of the <u>P</u>P2A Inhibitory <u>P</u>athway (PIP; Jak2-hnRNPA1-SET)^2^. Gain or loss of miRNA expression/function is a feature of cancer cells including CSCs^4, 5^ in which they may function as oncogenes or tumor suppressors influencing CSC’s function, tumor growth and innate anti-cancer immunity^6, 7^. Several miRNAs have been associated with CSC expansion and maintenance^4^ and PP2A inactivation^8^; however, a clear causal link between expression of specific miRNAs, persistence of the drug-resistant quiescent CSCs and PP2A loss-of-function is still missing. Furthermore, these miRNAs mostly act as regulators of CSC survival, maintenance into dormancy and cell cycle re-entry but it is unclear whether any of them directly promotes CSC cell cycle exit^4, 9–11^.

In myeloid leukemias, the mechanisms underlying LSC quiescence, survival and self-renewal, and reduced NK cell number and cytotoxicity result from integration of tumor cell-autonomous- and bone marrow (BM) microenvironment (BMM)-generated signals^12, 13^. The latter are triggered by BM niche-specific metabolic conditions (e.g. oxygen tension), cell-to-cell direct and, soluble and/or exosome-encapsulated factor (e.g. miRNAs, TGF*β*1)-mediated interactions between tumor, mesenchymal stromal (MSCs), endothelial (EC) and immune cells^6, 13–15^.

Among the miRNAs predicted *in silico* to re-activate PP2A in LSCs, we focused on hsa-*MIR300* (*MIR300*) because it was predicted to activate PP2A upon inhibiting multiple PIP components and PP2A-regulated factors essential for CSC maintenance, cancer development/progression and G1/S cell cycle transition. The human *MIR300* (*MIR300*) is an intergenic miRNA that belongs to the 14q32.31 *DLK1-DIO3* genomic-imprinted tumor suppressor miRNA cluster B^16, 17^; it was found involved in loss-of heterozygosity, inhibited in several tumor types with high mitotic index and during epithelial-to-mesenchymal transition (EMT), and associated with a CSC phenotype^18–22^.

By using Philadelphia-positive (Ph^+^) chronic myelogenous leukemia (CML) in chronic (CP) and blastic (BC) phase, and complex karyotype (CK) acute myeloid leukemia (AML), as paradigmatic examples of stem cell-derived neoplasms characterized by constitutive expression of oncogenic kinases (e.g. BCR-ABL1, JAK2), PP2A loss-of-function, altered miRNA expression and, impaired NK cell proliferation and cytotoxicity^2, 6, 23–27^, we show that *MIR300* is a BMM (hypoxia, MSC)-induced PP2A activator with cell context-independent tumor suppressor anti-proliferative and pro-apoptotic activities which are essential for induction and maintenance of LSC quiescence and impaired NK cell immunity but can also be exploited to selectively and efficiently induce cell death in qLSC and leukemic progenitor while sparing normal hematopoiesis.

## RESULTS

### Inhibition of *MIR300* tumor suppressor activities in leukemic progenitors and their dose-dependent differential induction in qLSCs

*MIR300* levels were progressively and markedly reduced by BCR-ABL1 activity (imatinib treatment) in BM CD34^+^ CML (CP and BC) and CK-AML progenitors compared to normal BM (NBM) cells and up to 800-fold higher in qLSCs (CD34^+^CFSE^max^) than dividing CD34^+^ CML (CP and BC) and CK-AML progenitors (Fig. 1a, b). Accordingly, *MIR300* expression was higher in HSC-enriched CD34^+^CD38^-^ than committed CD34^+^CD38^+^ CML (CP and BC) BM cells from patient samples at diagnosis, and comparable to those in NBM and umbilical cord blood (UCB) CD34^+^ cell fractions (Fig. 1a).

**Figure 1.**
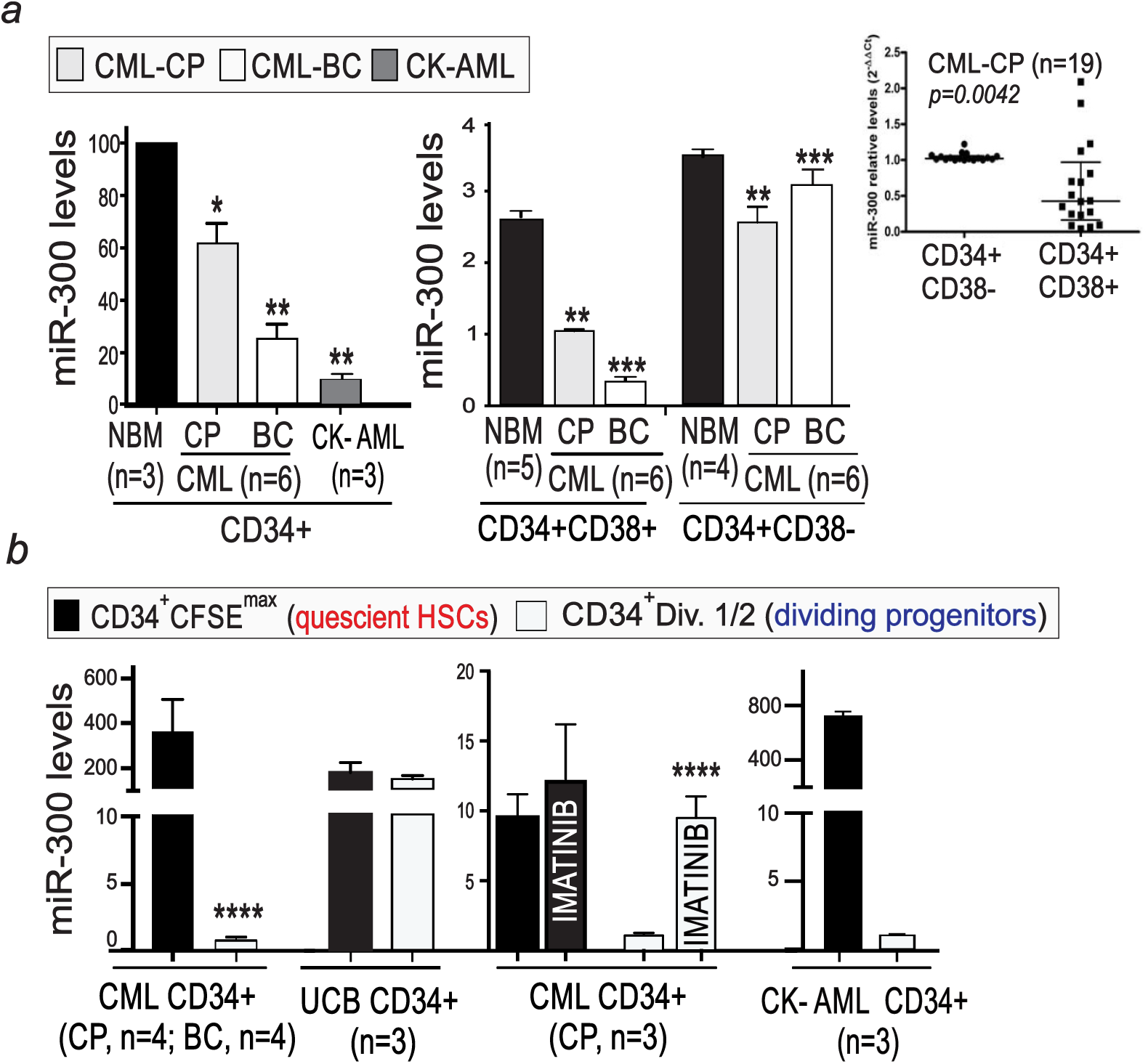
*MIR300* expression in quiescent leukemic stem and progenitor cells. (**a**) *(left) MIR300* levels in healthy (NBM) and CML CD34^+^ bone marrow (BM) cell fractions. *Inset* shows *MIR300* levels in additional CD38-fractionated CD34^+^ CML-CP BM cells expressed as n-fold difference in CD34^+^CD38^+^ compared to CD34^+^CD38^-^ samples. *(right) MIR300* levels in untreated and imatinib (IM; 24h)-treated CD34^+^ quiescent (CFSE^max^) and dividing (Div.1) CFSE-labeled CML, AML and umbilical cord blood (UCB) cells. Asterix on CD34^+^CD38^-^ cell populations (panel 1a) indicate significance between *MIR300* levels CD34^+^CD38^-^ vs CD34^+^CD38^+^ cells. Data are shown as mean ± SEM from at least three independent experiments. (∗*P*< 0.05, ∗∗*P* < 0.01, ∗∗∗ *P* < 0.001, ∗∗∗∗ *P* < 0.0001).

Consistent with the requirement of PP2A inhibition for leukemic but not normal stem/progenitor cell proliferation/survival^2, 28^, restoring *MIR300* expression at physiological levels by clinically-relevant *CpG-*based *MIR300* (*CpG-*miR-*300*) oligonucleotide (Fig. 2) or *MIR300*-lentiviruses (not shown) reduced by *≥*75% proliferation and clonogenic potential, and enhanced apoptosis (spontaneous and TKI-induced) of CD34^+^ CML (CP and BC) but not UCB cells (Fig. 2a, b). Importantly, Ki-67/DAPI (G0: *MIR300*≅46% vs. scr≅10%; G1: *MIR300*≅15% vs. scr≅50%; S/G2/M: *MIR300*≅4% vs. scr≅27.4%; and subG1: *MIR300*≅35% vs. scr≅3%) and FUCCI2BL-mediated (G1/G0: *MIR300*≅40.2% vs scr≅23.6%; G1/S: *MIR300*≅15% vs. scr≅2.85%; S/G2/M: *MIR300*≅45% vs. scr≅76.6%) cell cycle analyses in CD34^+^ CML (CP and BC) and LAMA-84 cells indicated that *MIR300* also selectively induced cell-cycle arrest with marked expansion of the G0 qLSC and sub-G1 apoptotic CD34^+^ cell fractions and reduced number of cells in G1 and S phases (Fig. 2c), indicating that *MIR300* functions as a dual activity (anti-proliferative and pro-apoptotic) tumor suppressor miRNA.

**Figure 2.**
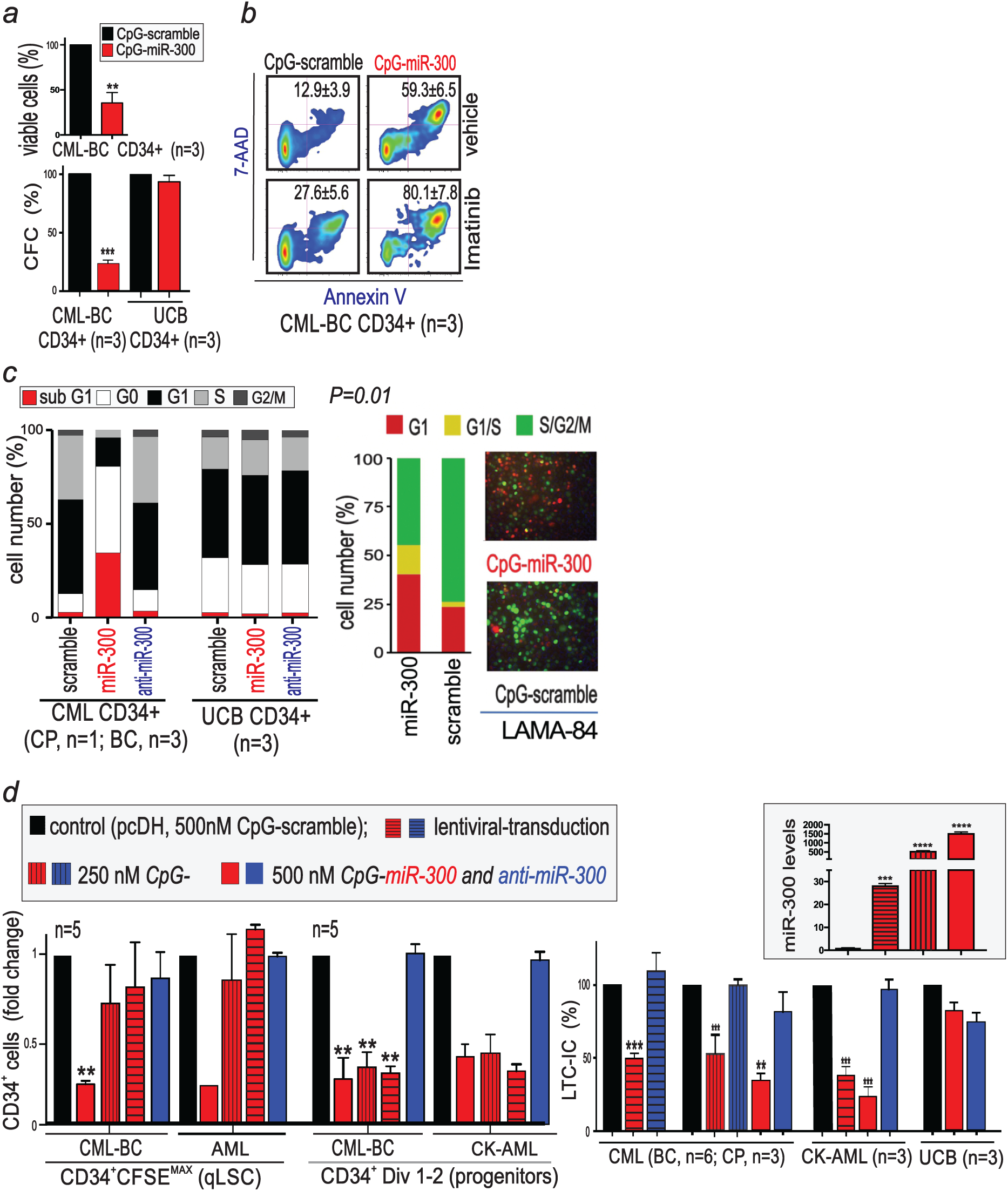
*MIR300* activity in quiescent leukemic stem and progenitor cells. (**a**) Growth (48h) and clonogenic potential (CFC) of *CpG-scramble-* and *CpG-MIR300*-treated (500 nM) CD34^+^ CML-BC and UCB cells. (**b**) Effect of *CpG-miR-300*or *CpG-scramble* (500 nM) on spontaneous and IM (18h)-induced apoptosis (Annexin V/7-AAD) in CD34^+^ CML-BC cells. (**c**) Ki-67/DAPI (left) and FUCCI2BL (right) cell cycle analysis of UBC and Ph^+^ (primary CD34^+^ and synchronized LAMA-84) cells exposed to the indicated *CpG-ON.* (**d**) Dose-dependent *MIR300* effect on: CML/AML qLSC (CFSE^max^) and progenitor (Div. 1-2) cell numbers (*left*) and on LTC-IC activity (*right*). Vector transduced and 500nM *CpG-scramble and -anti-MIR300* served as controls. *Inset: MIR300* levels in lentiviral-transduced and 250-500 nM CpG-*MIR300*-treated Ph^+^ cells. Data are shown as mean ± SEM from at least three independent experiments. (∗*P*< 0.05, ∗∗*P* < 0.01, ∗∗∗ *P* < 0.001, ∗∗∗∗ *P* < 0.0001).

The effects of *MIR300* modulation on quiescent stem cell number (CFSE assay), proliferation, survival and self-renewal (LTC-IC and CFC/replating) were assessed in CML (CP and BC) and CK-AML and healthy individuals (Fig. 2d). Inhibition of *MIR300* function (anti-miR-300) did not alter quiescent LSC and HSC number and activity (Fig. 2d, Supplementary Fig. 1). By contrast, graded *MIR300* expression (Fig. 2d, inset) differentially and selectively suppressed CML and AML but not UCB LTC-IC-driven *in vitro* hematopoiesis, LSC-derived clonogenic and self-renewal activity (Fig. 2d, *right* and Supplementary Fig 1), and quiescent (CFSE^max^) LSC numbers (Fig. 2d, *left*). Importantly, low-*MIR300* doses strongly inhibited CML and AML LTC-ICs and CFC/replating activities without affecting qLSC numbers whereas high-*MIR300* doses also reduced by ∼80% CML and AML qLSCs and dividing CD34^+^ progenitor cell numbers (Fig. 2d and Supplementary Fig 1). Thus, low *MIR300* expression may reduce colony-formation by impairing qLSC ability to proliferate and undergo cytokine-induced differentiation but have no effect on LSC survival that, instead, is halted at *MIR300* levels similar to those detected in qLSCs.

### *MIR300* acts as master PP2A activator and an inhibitor of G1/S transition

Gene ontology (GO) and Kyoto Encyclopedia of Genes and Genomes (KEGG) functional enrichment and clustering of *MIR300* predicted and validated mRNA targets (DIANA-mirPath v.3) indicated that *MIR300*-induced massive apoptosis of qLSCs and leukemic progenitors likely depends on PP2A activation triggered by *MIR300*-induced inhibition of SET and other PIP factors (Fig. 3a). Likewise, *MIR300* targeting of G1/S cell cycle regulators CCND2 (cyclin D2) and CDK6 and of several other PP2A-regulated survival and mitogenic factors (e.g. CTNNB1, JAK2, MYC, Twist1) (Fig. 3a, and Supplementary Fig. 2a *left*, 2b) may contribute to *MIR300*-induced cell death and account for cell cycle arrest and expansion of the qLSC G0 compartment.

**Figure. 3.**
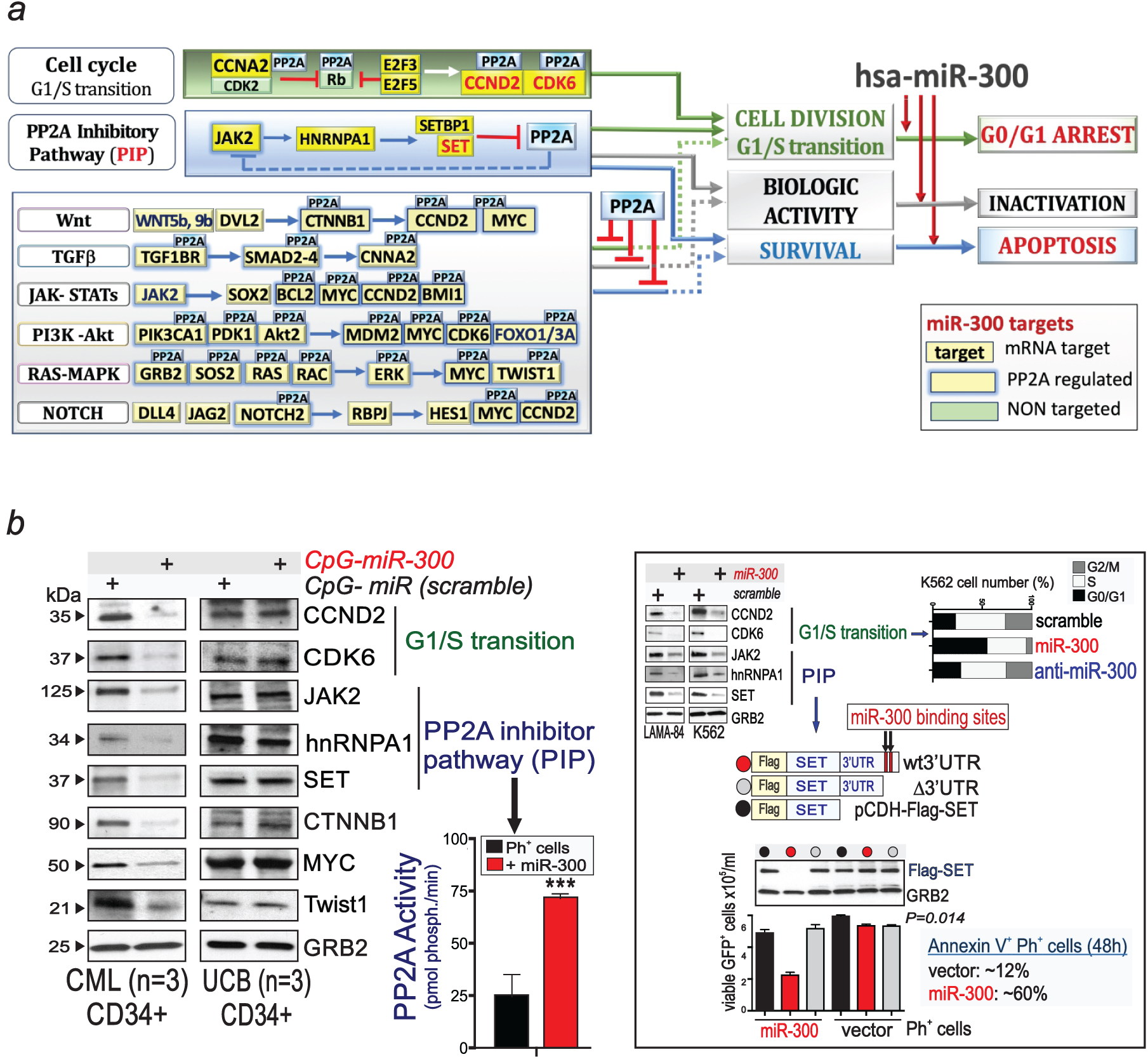
*MIR300* is a master PP2A activator and an inhibitor of G1/S transition. (a) KEGG/GO analysis of the effect of *MIR300* on PP2A-regulated signal transduction pathways. **(b)** *Left*: Representative blots show effect of *MIR300* on its targets and PP2A activity in UCB and CML-BC CD34^+^ cells and cell lines exposed to *CpG-scramble* and *CpG-miR-300*(500 nM; 48-72h). *Right*: (top) Dapi/Ki67 cell cycle analysis of *CpG-scramble, -MIR300 and -anti-MIR300* (500 nM; 21h)-treated aphidicolin-synchronized K562 cells; (middle) Flag-SET lentiviral constructs with wild type or a deleted mRNA 3′UTR; (bottom) *MIR300*-induced downregulation of Flag-SET proteins, and rescue of Ph^+^ cells from exogenous *MIR300*-induced growth inhibition (trypan blue exclusion)/apoptosis (Annexin V^+^) by Flag-SET cDNAs lacking *MIR300* binding site. Similar results were obtained with LAMA-84 cells (not shown).

Indeed, restoring *MIR300* expression in primary CD34^+^ CML (CP and BC) progenitors and Ph^+^ cell lines by 500nM *CpG-miR-300*treatment, re-activated PP2A by inhibiting SET and the other PIP molecules (i.e. JAK2, hnRNPA1) and markedly reduced CDK6, CCND2, CTNNB1 (*β*-catenin), Twist1 and MYC expression (Fig. 3b). Note that CTNNB1, CCND1/2 and Twist1 are validated *MIR300* targets^20, 21^. Furthermore, consistent with the notion that high levels of active PP2A are not detrimental in normal CD34^+^ quiescent stem and committed progenitors^2, 28^, *CpG-miR-300* did not alter levels of *MIR300* targets in CD34^+^ UCB cells (Fig. 3b). Importantly, expression of *MIR300*-insensitive Flag-tagged *SET* mRNAs deleted of the whole 3’UTR (Flag-SET) or just a region encompassing the high- and low-affinity *MIR300*-binding sites (Flag-*Δ*3’UTR-SET) but not that of full-length wild type SET mRNA (Flag-wt3’UTR-SET), rescued Ph^+^ progenitors from *MIR300*-induced cell cycle arrest and PP2A-dependent apoptosis (Annexin V^+^ cells) (Fig. 3b, *right*).

Interestingly, hierarchically clustering of *MIR300* predicted and validated targets based on integration of algorithms (DIANA microT-CDS, mirDIP 4.1, ComiR and CSmiRTar) that also take into account *MIR300* expression levels in normal and leukemic BM cells, positioned the G1/S cell cycle regulators CCND2 and CDK6 within the top 2% and SET within the 5% of *MIR300* targets (Fig. 4a). Conversely, JAK2, hnRNPA1, CTNNB1 and Twist1 fell within the top 1/3, and Myc in the lower 2/3 of *MIR300*-interacting mRNA cluster (Fig. 4a, and Supplementary Fig. 3). Notably, other factors contributing to G1/S transition (e.g. CNNA2) and LSC re-entry into cycle for self-renewal or maturation (e.g. Notch signaling) clustered together with CCND2 and CDK6 (Supplementary Fig. 3). By contrast, *MIR300* targets belonging to JAK-STATs, PI3K-Akt, Wnt, MAPK signaling pathways, which regulate survival and expansion of leukemic progenitors and disease progression, ranked below SET and within the bottom 75% of *MIR300* target distribution (Supplementary Fig. 3). Because most of these *MIR300* targets are also validated target of PP2A enzymatic activity (Supplementary Fig. 2b, 3), their PP2A-dependent inactivation likely occurs upon *MIR300*-induced SET inhibition. Thus, MIR-300-dependent inhibition of CCND2 and CDK6 should precede that of SET which should be followed by the PP2A-dependent JAK2, CTNNB2, TWIST and, lastly, MYC inactivation.

**Figure. 4.**
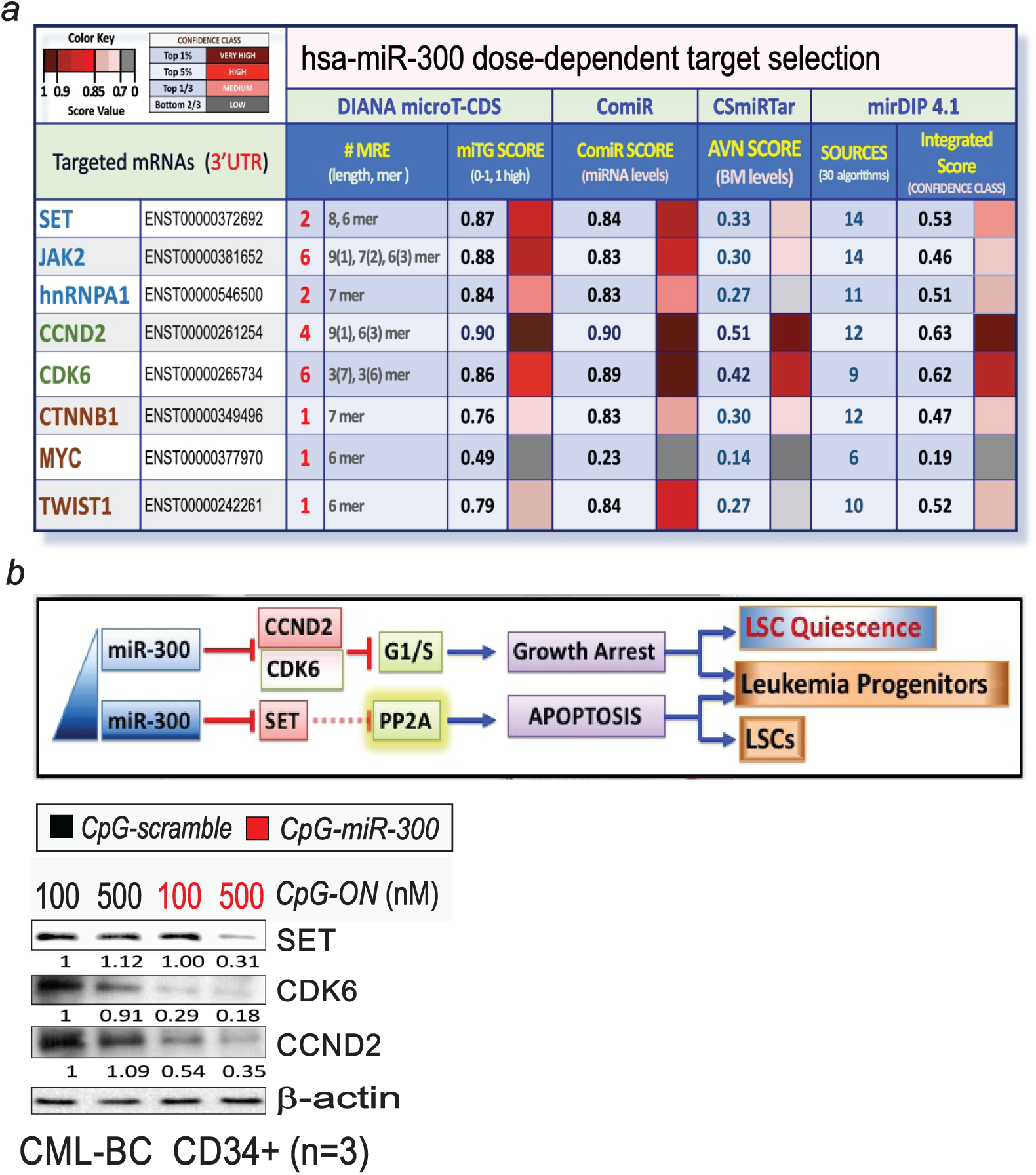
*MIR300* dose-dependent mRNA target selection. (a) Hierarchical clustering of statistically significant (P<0.05 with FDR correction) *MIR300* targets using the indicated databases (number of binding sites is indicated in red). **(d)** (top) Biological Effects of *MIR300* dose-dependent target selection mechanisms; (bottom) SET, CDK6, CCND2 and *β*-actin levels in *CpG-MIR300*- and *CpG*-scramble-treated (100-500 nM; 48h) CML-BC CD34^+^ cells.

Indeed, immunoblots confirmed that CCND2 and CDK6 inhibition occurs in CD34^+^ leukemic stem/progenitor cells at a *CpG-miR-300* concentration (100 nM) not triggering qLSC apoptosis whereas SET inhibition requires the same *CpG-miR-300* dose that reduces qLSC numbers (Fig. 4b). This is consistent with the presence of 4 (1 very high affinity 9mer), 6 (3 high affinity 7mer) and 2 (1 high affinity 8mer) *MIR300* binding sites in *CCND2, CDK6* and *SET* mRNA 3’UTRs, respectively (Fig. 4a). Because CDK6/CCND2 downregulation is an essential feature of G1/G0-arrested qLSCs^29, 30^ and SET inhibition is sufficient for inducing PP2A-dependent LSC and progenitor cells apoptosis^2, 26^, the ability of *MIR300* to sequentially trigger growth arrest and PP2A-mediated apoptosis of CD34^+^ CML/AML LSC and progenitors (Fig. 2c, d) indicates that *MIR300* tumor suppressor activities are cell context-independent and suggests that *MIR300* expression in LSC may account for their entry into quiescence whereas its downregulation in leukemic progenitors likely occurs to prevent apoptosis (Fig. 4b, top).

Specificity of *MIR300* activity is further supported by the inability of intragenic 14q32.31 tumor suppressor^17^ hsa-miR-381-3p (miR-381-3p) to downregulate JAK2, SET and *β*-catenin levels and inhibit CD34^+^ CML cell proliferation and clonogenic potential (CFC assays) despite having an identical seed sequence, highly homologous flanking region and similar expression patterns in LSC-enriched CD34^+^CD38^-^, committed CD34^+^CD38^+^ progenitors and bulk CD34^+^ CML-CP and - BC cells (Supplementary Fig. 4a). Note that, the presence of a +4 A-to-G seed sequence mutation in the mouse *mir300* (mmu-miR-300) gene significantly alters target repertoire (miRTar, MFE: *≤* - 10kcal/mol; Score: *≥*136.5) and it is predicted to impair mmu-miR-300 tumor suppressor activities by inhibiting binding to *JAK2, SET, CCND2 and CDK6* mRNAs (not shown).

### BMM-induced C/EBPβ-dependent *MIR300* tumor suppressor activity preserves LSCs

*MIR300* is under the control of an intergenic differentially-methylated region (IG-DMR) preceding and controlling the expression of MEG3 (Supplementary Fig. 5a), a maternally-expressed genomic imprinted anti-proliferative lncRNA that is important for long-term HSC (LT-HSC) maintenance but strongly inhibited upon promoter methylation in virtually all types of cancer including CD34^+^ CML cells^17, 31^. Treatment with 5-Aza-2’-deoxycytidine (5-Aza) augmented by 10^4^-10^5^-fold *MIR300* expression in Ph^+^ cells (Fig. 5a, *left*); however, nearly all cells underwent apoptosis after 24h (not shown), suggesting that *MIR300* upregulation in qLSCs unlikely depends on IG-DMR demethylation and expression of all 74 cluster B tumor suppressor miRNAs (Supplementary Fig. 5a). Thus, *MIR300* induction in CML/AML qLSCs and its ability to increase the quiescent fraction of leukemic but not normal CD34^+^ cells implies that BM osteogenic niche metabolic and cellular factors (e.g. MSCs, hypoxia), known to inhibit growth and induce quiescence of leukemic cells^12–14^, may selectively favor LSC entry into quiescence by increasing *MIR300* expression and regulating its anti-proliferative activity in CD34^+^ LSCs (Fig. 5a *right*).

**Figure 5.**
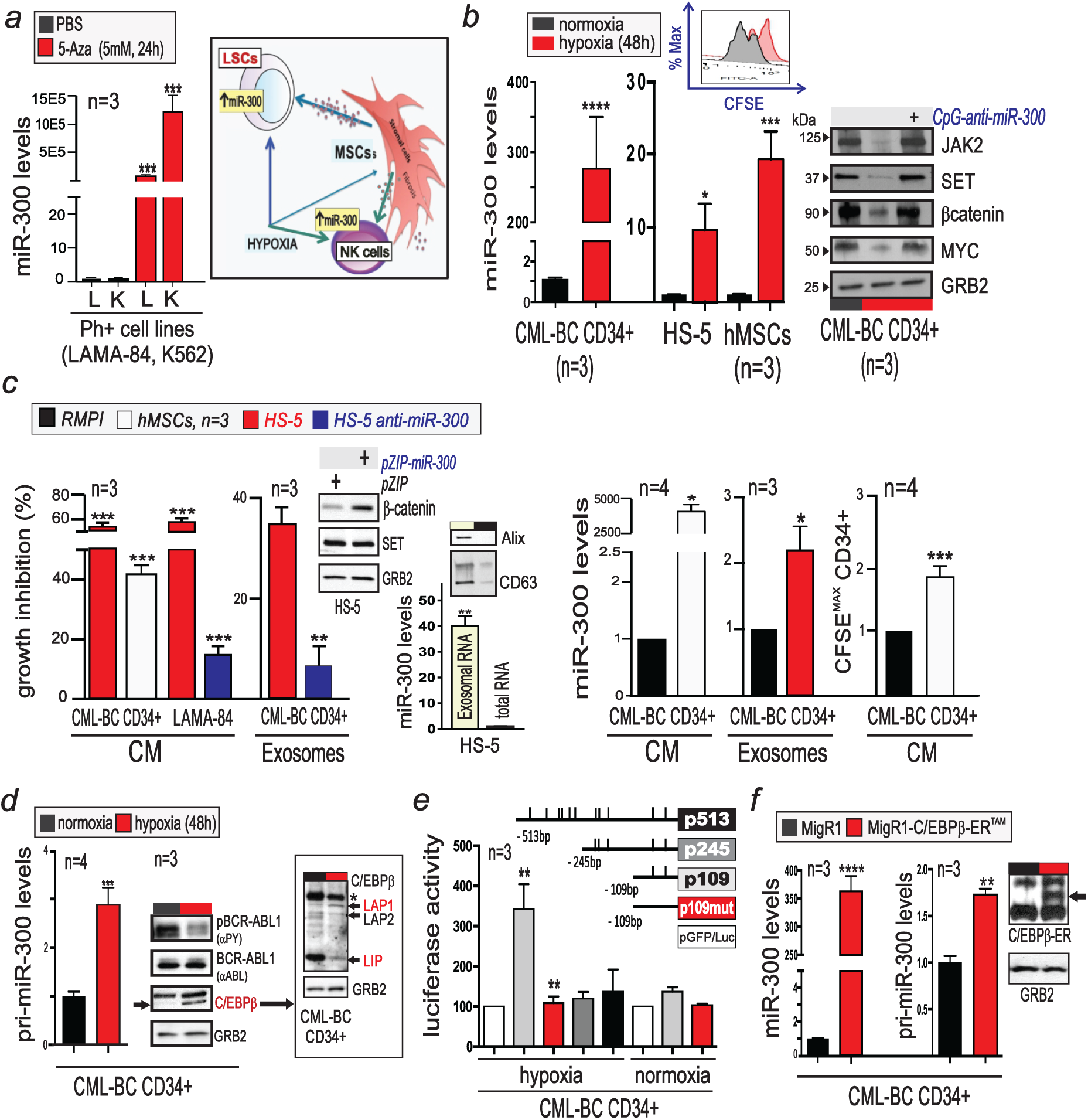
BMM-induced C/EBPβ-dependent *MIR300* tumor suppressor activity preserves LSCs. (a) *left*: *MIR300* levels in 5-Aza- or DMSO-treated (24h) Ph^+^ cells. *right*: *MIR300* regulation by the BMM in LSCs and NK cells. **(b)** Effect of hypoxia on: i) *MIR300* levels in CD34^+^ CML-BC, BM-derived primary (hMSCs) and HS-5 MSCs; and ii) *MIR300* targets in untreated and *CpG-anti-MIR300*-treated (500 nM, 48h) and CML-BC cells. Inset: effect of hypoxia on CFSE^+^CD34^+^ CML-BC proliferation. **(c)** Effect of hMSC- and HS-5-CM and/or exosomes (50-100 μg/ml) from parental, vector (pZIP) and anti-*MIR300* (pZIP-*MIR300*)-transduced primary hMSCs and/or HS-5 cells on: i) proliferation (% growth inhibition); ii) qLSC fraction (CFSE^max^CD34^+^); and iii) *MIR300* levels in CD34^+^ CML-BC and LAMA-84 cells. Insets: *left*, β-catenin and SET levels in anti-*MIR300* (*pZip-300*)-transduced HS-5; *right*, *MIR300* in HS-5 Alix^+^CD63^+^ exosomes. **(d)** Effect of hypoxia on primary *MIR300* transcripts (pri-*MIR300*), C/EBP*β* (LAP1, LAP2 and LIP isoforms), BCR-ABL1 expression (*α*ABL) and activity (*α*PY), and GRB2 levels in CD34^+^ CML-BC cells. (*): nonspecific band. **(e)** *MIR300* promoter/enhancer activity in hypoxia-(48h) and normoxia-cultured CML-BC CD34^+^ cells transduced with pGFP/Luc-based *MIR300*-reported constructs. p109mut is mutated in the -64 and -46 bp C/EBP*β* binding sites. **(f)** Effect of ectopic C/EBP*β* (*inset*) on mature (*MIR300*) and primary (pri-*MIR300*) *MIR300* levels in CD34^+^ CML-BC cells. Data are represented as mean ± SEM for at least three experiments.

Indeed, expression of *MIR300*, but not of miR-381-3p (Supplementary Fig. 4b), increased at qLSC or higher levels (Fig. 1b) in primary CD34^+^ CML-BC and/or LAMA-84 cells exposed to hypoxia, BM-derived primary CD34^-^CD45^-^CD73^+^CD105^+^CD90^+^CD44^+^ hMSC and HS-5 conditioned medium (CM) or *MIR300*-containing CD63^+^Alix^+^ MSC exosomes (Fig. 5b,c). This correlated with a 40-60% reduced cell division and, importantly, with doubled numbers of CD34^+^ CML cells in the CFSE^max^ CD34^+^ qLSC fraction (Fig. 5b, c and Supplementary Fig. 5b). Similarly, Ph^+^ LAMA-84 (Supplementary Fig. 5c) and K562 (not shown) cells responded to MSC-CM exposure by slowly ceasing proliferation and modifying gene expression in a manner similar to that of CML qLSCs^2, 12^. In fact, exposure to MSC (CM and/or exosomes) neither altered SET and PP2Ac^Y307^ (inactive) levels (Supplementary Fig. 5c) nor induced cell-death (not shown) in CD34^+^ CML-BC and/or Ph^+^ cells, suggesting that MSCs increase *MIR300* expression at levels not sufficient to trigger PP2A-mediated apoptosis.

The absolute requirement of *MIR300* for BMM-induced LSC quiescence is revealed by the ability of anti-*MIR300* molecules (pZIP-miR-300 or CpG-anti-miR-300) to prevent hypoxia-induced downregulation of *MIR300* targets (e.g. JAK2, Myc, SET and *β*-catenin) in CD34^+^ leukemic cells, and to protect CD34^+^ CML cells from MSC (CM and exosome) growth-inhibitory activity when expressed in MSCs, in which they enhance *β*-catenin but not SET expression (Fig. 5b, c). Moreover, the evidence that apoptosis is not induced in MSC-exposed CD34^+^ CML cells despite the marked *MIR300* increase, and that hypoxia but not MSC CM induces *MIR300*-dependent SET inhibition and increases mature *MIR300* to levels similar to those in qLSCs (Fig. 1b and 5b) indicate that the contribution of MSC (CM and exosomes) to *MIR300* expression in LSCs is not sufficient for inhibiting SET and triggering PP2A-dependent apoptosis, and that hypoxia may induce *MIR300* transcription in LSCs. Accordingly, primary *MIR300* (pri-miR-300) transcripts were substantially higher in CD34^+^ CML-BC cells cultured (n=4; 48h) in hypoxic than normoxic conditions (Fig. 5d).

Luciferase (luc) assays in normoxia- and hypoxia-cultured CD34^+^ CML-BC cells transduced with reporter constructs containing the full-length (p513) or a 5’-deleted (p245, p109) intergenic region, identified a hypoxia-sensitive *MIR300* regulatory element within the 109 bp preceding the human *MIR300* gene (Fig. 5e). Because ENCODE (V3) ChIP-Seq and PROMO analyses revealed that the CCAAT Enhancer Binding Protein B (C/EBP*β*) may interact with two DNaseI hypersensitive regions within the intergenic 513bp preceding the *MIR300* gene (Fig. 5e), we assessed whether hypoxia-induced *MIR300* upregulation results from C/EBP*β* binding and transactivation of *MIR300* transcription.

Luciferase assays with a p109 construct carrying mutated C/EBP*β* binding sites at position -64 and -46 bp (p109mut) indicate that the integrity of these hypoxia-sensitive C/EBP*β* responsive elements is essential for *MIR300* transactivation in CD34^+^ CML-BC cells (Fig. 5e). Indeed, increased pri-miR-300 expression in hypoxic CD34^+^ CML-BC cells correlated with markedly higher C/EBP*β* LAP1 (transcriptionally active) levels (Fig. 5d). By contrast, lack of C/EBP*β*-driven p109-luc activity in normoxic CD34^+^ CML-BC progenitors correlated with increased C/EBP*β* LIP (inhibitory function) expression (Fig. 5d), indicating that hypoxia-induced *MIR300* expression requires C/EBP*β* transactivation. Importantly, consistent with the notion that BCR-ABL1 is inactivated by hypoxia to induce quiescence and allow survival of CML LSCs^2, 14^ and that BCR-ABL1 activity inhibits *CEBPB* translation^32^, *MIR300* induction by hypoxia correlated with decreased BCR-ABL1 activity but not expression (Fig. 5d), and ectopic C/EBP*β*-ER^TAM^ mimicked the effect of hypoxia on C/EBP*β* by rescuing primary (pri-miR-300) and mature *MIR300* expression in normoxic CD34^+^ CML-BC progenitors (Fig. 5f).

Importantly, the notion that mouse c/ebp*β* induces BCR-ABL^+^ LSK exhaustion^33^ does not argue against a role for human C/EBP*β* as an inducer of LSC quiescence because a) LSKs are pushed into cycle by a fully active BCR-ABL1 that promotes C/EBP*β*-dependent LSK maturation^33^; b) the hypoxia-sensitive human C/EBP*β*-responsive *MIR300* regulatory element is not conserved in mouse cells; and, c) an A-to-G mutation in the mmu-miR-300 seed sequence (Supplementary Fig. 4) prevents mmus-miR-300 targeting of *ccnd2* and *cdk6* mRNAs (not shown).

Hypoxia also substantially increased by 10-20-fold *MIR300* and C/EBP*β* protein but not mRNA levels in BM-derived primary MSCs and/or HS-5 cells (Fig. 5b and Supplementary Fig. 5d), suggesting that the hypoxic conditions of the osteogenic BM niche may simultaneously induce C/EBP*β*-dependent *MIR300* transcription in LSCs and increase the amount of MSC-derived exosomal *MIR300* transferred into LSCs. Conversely, *MIR300* levels in C/EBP*β−*responsive CD34^+^ CML-BC cells and/or Ph^+^ cell lines were not influenced by ectopic C/EBP*α* expression or exposure to TGF*β*1 blocking-Ab (Supplementary Fig. 5e, f).

Thus, increased *MIR300* transcription occurs in a TGF*β*-independent manner through the hypoxia-induced C/EBP*β* LAP1 expression in LSCs and MSCs. The latter also contribute to increased *MIR300* levels in LSCs via exosomal transfer.

### BMM-induced MIR300 inhibits NK cell anti-cancer immunity

Cytokine-induced expression of miR-155 and SET, and inhibition of PP2A activity (Fig. 6a and Supplementary Fig. 6a) are essential for CD56^+^CD3^-^ primary NK and clinically-relevant^34^ NK-92 cell proliferation and cytotoxicity against tumor-initiating cells^35^ including CD34^+^ CML qLSCs (Fig. 6b, *left*). Conversely, endosteal BMM factors elicit signals that inhibit NK cell proliferation and cytotoxicity^7, 35, 36^ and are, likely, responsible for dysfunctional NK cells in leukemia patients^24^. Thus, we investigated whether BMM-induced *MIR300* inhibits NK cell killing of CML qLSCs and progenitors through downregulation of key regulators of NK cell proliferation (e.g. CCND2, CDK6) and activity (e.g. SET, miR-155) (Fig. 6a). First, we found that increased *MIR300* expression is also a feature of peripheral blood (PB) CD56^+^CD3^-^ NK cells isolated from CML patients at diagnosis but not of NK cells from healthy individuals (Fig. 6b). Furthermore, KEGG/GO analyses (Supplementary Fig. 2a) indicated that *MIR300* may reduce levels of key regulators of NK cell function and account for suppression of innate anti-cancer immunity in CML.

**Figure 6.**
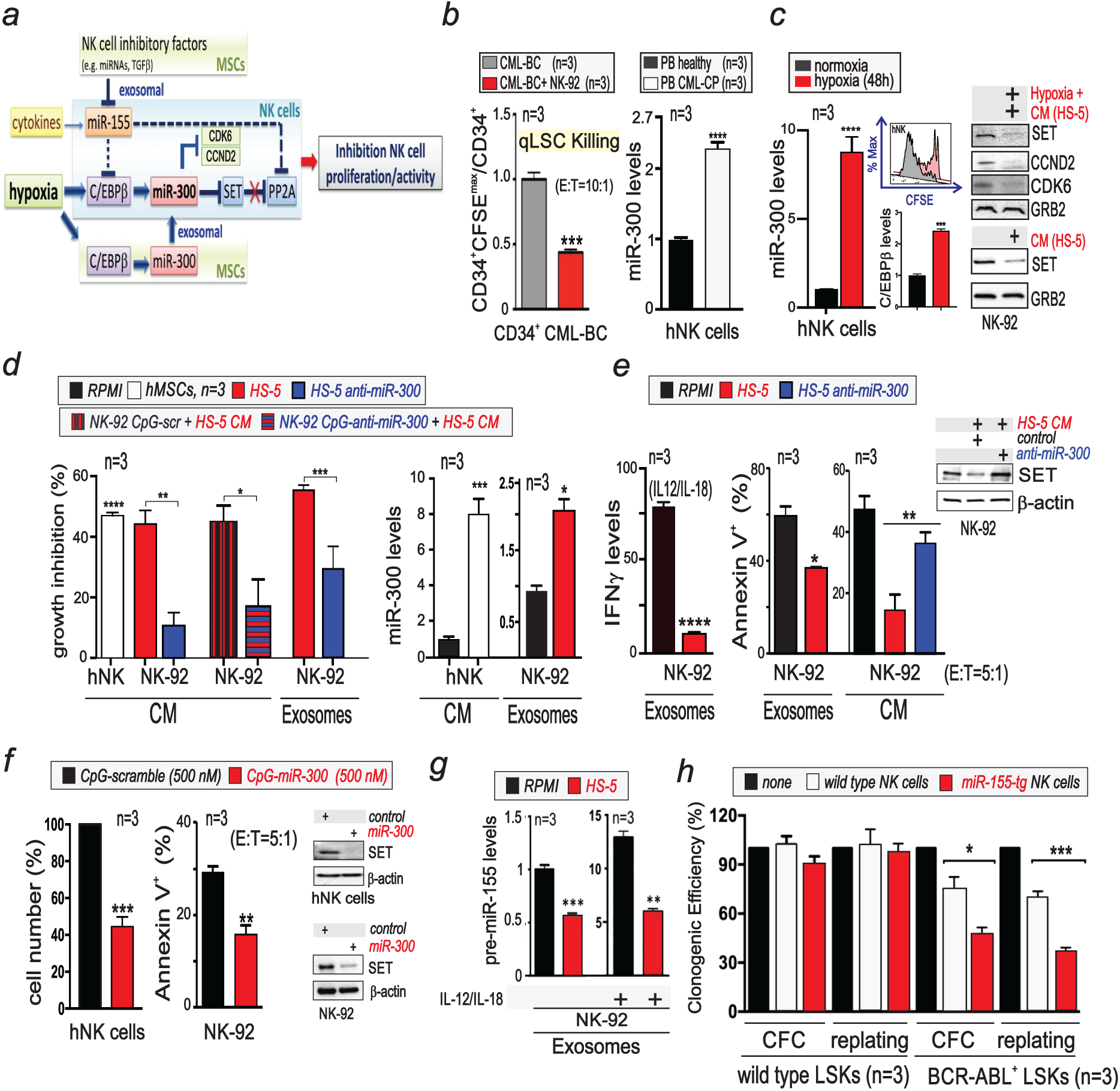
*MIR300* inhibits NK cell growth and activities. **(a)** BMM-induced *MIR300*/miR-155 interplay on NK cell proliferation/ activity. **(b)** *Left:* NK-92 cytotoxicity against CML-BC qLSCs. *Right: MIR300* levels in CD56^+^CD3^-^ NK (hNK) cells from healthy and CML-CP individuals. **(c)** Effect of hypoxia and MSC CM on *MIR300*, C/EBP*β*, SET, CCND2, CDK6 and GRB2 levels in hNK and/or NK-92 cells. *Inset:* effect of hypoxia on CFSE^+^ NK cell growth. **(d)** Effect of CM and exosomes from hMSCs and from parental, vector- and anti-*MIR300-transduced* and HS-5 cells on proliferation (*left*) and *MIR300* levels (*right*) in hNK and, untreated, 500 nM *CpG-scramble* or *CpG-anti-MIR300* treated NK-92 cells. **(e)** IL-12/IL-18 (18h)-induced IFN-γ mRNA levels and NK cytotoxicity (% Annexin V^+^ K562 cells) in IL-2-cultured NK-92 cells exposed to CM or exosomes from parental or vector- and anti-*MIR300*-transduced HS-5 cells. (Inset) SET and Actin levels in NK-92 exposed to CM from vector and anti-miR300 transduced HS-5 cells. RPMI medium served as control. **(f)** Effect of *CpG-scramble* or *CpG-miR-300*treatment (500 nM; 36h and 7d) on IL-2-depleted NK-92 cytotoxicity (% Annexin V^+^ K562 cells), IL-2-induced hNK cell proliferation (% cell number) and on SET and *β*-actin expression. **(g)** Precursor (pr*e-miR-155*) miR-155 (BIC) levels in resting and IL-12/IL-18 (18h)-stimulated NK cells exposed (48h) to HS-5 exosomes. **(h)** Cytotoxicity (18h; E:T 20:1) of wild type (wt) and miR-155 transgenic (miR-155-tg) NK cells against LSK cells from not-induced and leukemic SCL-tTA-BCR-ABL1 mice by CFSE^+^ LSK-driven methylcellulose CFC/replating assays and expressed as % changes in clonogenic potential of leukemic progenitors (CFC) and self-renewing LSCs (replating). Data are represented as mean ± SEM for at least three experiments.

Second, hypoxia (1% O_2_ tension) and/or BM MSC- and HS-5-derived CM and Alix^+^CD63^+^exosomes (100ug/ml) markedly increased C/EBP*β* and *MIR300* but reduced CCND2, CDK6 and SET expression (Fig. 6c). Consistent with the notion that downregulation of CCND2, CDK6 and SET^3, 36, 37^ impairs NK cell cytokine-dependent proliferation and induces PP2A-dependent NK cell inactivation, respectively, BMM (hypoxia and MSCs)-induced *MIR300* upregulation was accompanied by 45% to 75% inhibition of IL-2-dependent CD56^+^CD3^-^ primary NK and NK-92 cell proliferation (Fig. 6d and Supplementary Fig. 6b) and by severely impaired NK cell immunoregulatory (IFN*γ* production) and anti-tumor cytotoxic (K562 killing) activities (Fig. 6e). Importantly, lentiviral anti-miR-300 (pZIP-miR-300)-transduction into HS-5 cells (Fig. 6d, *left*) significantly suppressed MSC CM- and/or exosome inhibitory effects om NK cell proliferation and anti-cancer cytotoxicity (Fig. 6d, e). Similar effect was obtained by pre-treatment of NK-92 cells with *CpG-anti-miR-300* but not with *CpG-scramble* (Fig. 6d). This suggests that BMM-induced impaired NK cell growth and anti-qLSC cytotoxicity occurs in a *MIR300*-dependent manner. In fact, *CpG-miR-300* treatment mimicked the inhibitory effects of exposure to BMM (hypoxia and MSCs) and strongly suppressed IL-2-induced proliferation and cytotoxicity of CD56^+^CD3^-^ NK and NK-92 cells (Fig. 6f). However, sequencing (RNAseq) of MSC exosomal RNA suggested that also other MSC-derived miRNAs may potentially impair NK cell activity (Supplementary Fig. 6c). Interestingly, MSC HS-5 exosomes also reduced miR-155 (BIC) transcription in IL-12/IL-18-stimulated and resting NK-92 cells (Fig. 6g). Because, miR-155 not only inhibits SHIP1 and PP2A to allow MAPK- and AKT-dependent NK cell proliferation and cytotoxic activity^8, 35, 38^ but also suppresses C/EBP*β* expression^39^, MSC-induced transcriptional downregulation of miR-155 precursor levels likely contributed to C/EBP*β*-mediated induction of *MIR300* (Fig. 6a, g).

To determine whether constitutive miR-155 overexpression in NK cell enhances their anti-LSC cytotoxic activity, we employed highly proliferating lck-miR-155 (miR155-tg) transgenic NK1.1^+^CD3^-^ NK cells, which show enhanced proliferation and cytotoxic activity in vivo^35^. miR155-tg NK1.1^+^CD3^-^ NK cells killed self-renewing CFSE^+^ leukemic SCL-tTA-BCR-ABL1 but not normal LSC-enriched LSK cells more efficiently than wild type (wt) NK1.1^+^CD3^-^ NK cells (∼70% vs ∼30% cytotoxic activity) (Fig. 6h), suggesting that therapies with allogeneic NK cells engineered to express high miR-155 levels alone or associated with *MIR300* downregulation may overcome BMM-induced loss of NK cell proliferation/activity. Indeed, clinically-relevant^31, 40–44^ NK-92 cells already decreased by 50% TKI-resistant CML-BC qLSC numbers (Fig. 6b).

### TUG1 selectively allows MIR300 anti-proliferative activity in CML and AML LSCs

Acute and chronic myeloid qLSCs display low to undetectable CCND2 and CDK6 expression but high SET (PP2A inhibition) levels^2, 23, 30^, and survive despite the high levels of functional (target downregulation) *MIR300* (Fig. 1b, 5b), suggesting that there must be a *MIR300*-interactinge factor that uncouples and differentially regulates *MIR300* activities to allow LSC cell cycle exit and prevents qLSC apoptosis through modulation of free *MIR300* intracellular levels. In this regard, the taurine upregulated gene 1 (TUG1) was described as a *MIR300-*interacting lncRNA in gallbladder carcinoma and enhances tumor cell growth and EMT upon binding to miRNAs and preventing suppression of a set of mRNAs (e.g. Myc, CCND1/2, CTNNB1, Twist1) regulating stemness and described as *MIR300* target in solid tumor cells^22, 45, 46^, suggesting that TUG1 may differentially regulate *MIR300* tumor suppressor activities (Fig. 7a).

**Figure 7.**
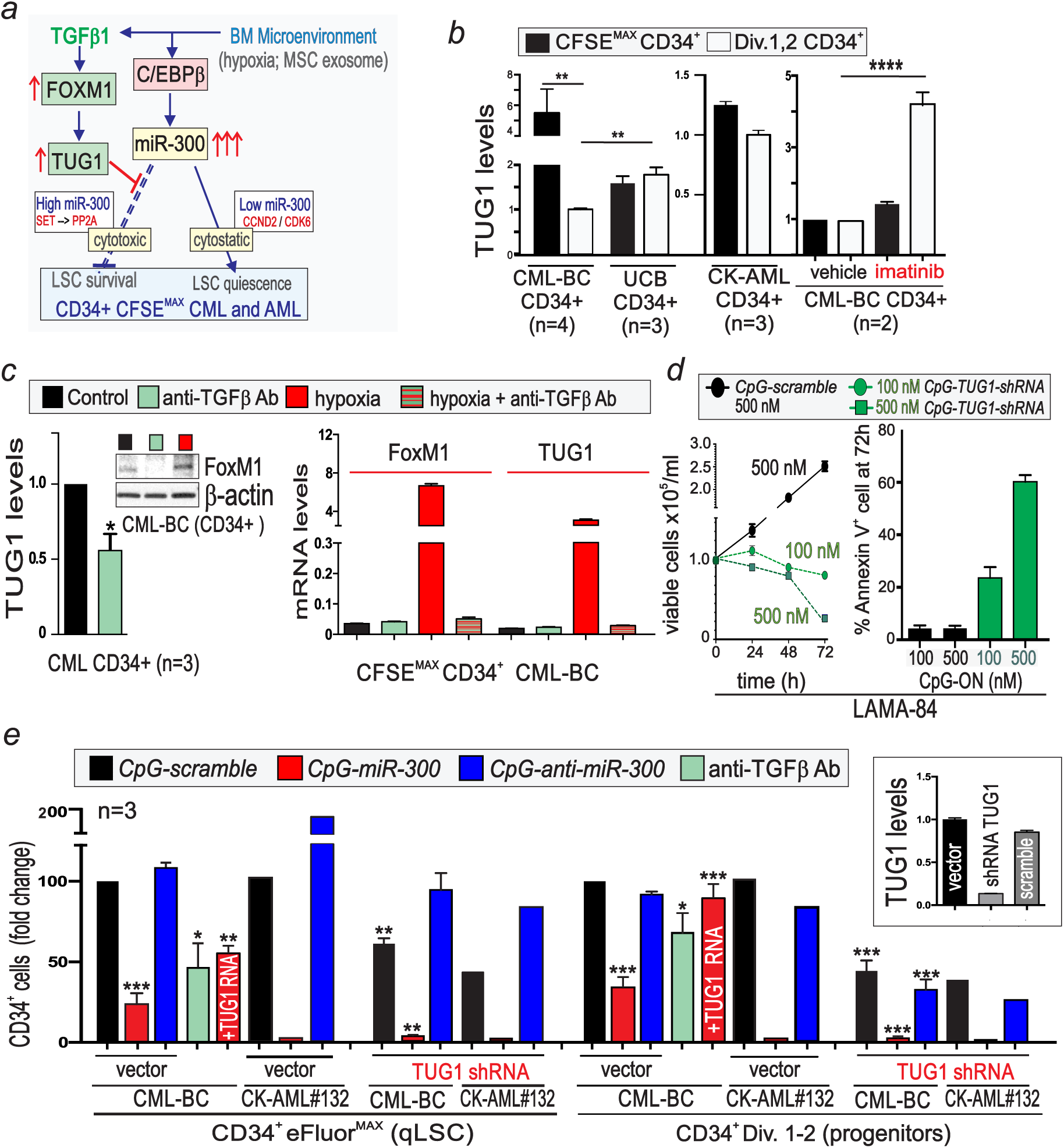
*TUG1 selectively allows MIR300 anti-proliferative activity in LSCs*. (a) *MIR300*-TUG1 interplay regulating LSC survival and quiescence. **(b)** TUG1 levels in CD34^+^ quiescent stem (CFSE^max^) and dividing progenitors (Div.1,2) and in untreated and imatinib-treated CD34^+^ cells **(c)** Effect of anti-TGF*β* antibody (Ab) on TUG1 lncRNA and FoxM1 levels in CD34^+^ (*left)* and CD34^+^CFSE^max^ CML-BC cells exposed (48h) to 1% O_2_. **(d)** Effect of anti-TGF*β* Ab, TUG1 shRNA, TUG1 RNA and control (CpG-scramble or empty vector) on recovery of untreated and *CpG-scramble*, *-MIR300* and/or *-anti-MIR300* eFluor^+^CD34^+^ CML/AML qLSCs (eFluor^max^) and dividing (Div.1-2) progenitors relative to input. *Inset*: TUG1 levels in vector, *TUG1 shRNA* and scramble-shRNA cells. Data are represented as mean ± SEM for at least three experiments.

Accordingly, TUG1 levels are markedly increased in CD34^+^ normal (UCB and BM) and leukemic HSCs including CD34^+^ qLSCs from CML-BC and CK-AML patients and healthy individuals (Fig. 7b and Supplementary Fig. 7b, e). In CML, TUG1 is regulated in a manner similar to that of *MIR300* (Fig. 1, 7b); in fact, TUG1 expression in qLSCs is imatinib-insensitive whereas it is markedly reduced by BCR-ABL1 activity in dividing CD34^+^ progenitors (Fig. 7b). In AML, TUG1 expression is induced by AML oncogenes (e.g. AML1-ETO, MLL-fusions) and readily detectable in different AML subtypes (Supplementary Fig. 7b, c). Interestingly, TUG1 levels were not downregulated in dividing CD34^+^ CK-AML progenitors but remained similar to those detected in CK-AML qLSCs (Fig. 7b).

Because TUG1 expression correlates with that of TGF*β*1 and TGFBR1 in tumor cells undergoing EMT^47^ and TGF*β*1 is known to be important for LSC quiescence and secreted by leukemic cells including CD34^+^ CML blasts^48^, we investigated the role of TGF*β*1 in TUG1 upregulation and *MIR300*-induced growth-arrest of CD34^+^ leukemic cells and qLSC apoptosis (Fig. 2, 5). Exposure (48h) of CD34^+^ CML cells to a TGF*β*1 blocking antibody (anti-TGF*β* Ab; *green bars*) halves TUG1 levels and qLSC numbers but very modestly affects dividing leukemic progenitors (Fig. 7c *left*, 7e). Interestingly, exposure to anti-TGF*β* Ab also reduces levels of FoxM1 (Fig. 7c), a transcription factor that mediates HSC quiescence and induces TUG1 expression in osteosarcoma cells^49, 50^. By contrast, FoxM1 and/or TUG1 expression is strongly induced in a TGF*β*-dependent manner in hypoxia-cultured (48h, 1% O_2_) CML CD34^+^CFSE^max^ qLSCs and bulk CD34^+^ CML-BC cells (Fig. 7c), suggesting that hypoxia-induced TGF*β*1^48^ may enhance TUG1 levels in a FoxM1-dependent manner to neutralize *MIR300* pro-apoptotic activity in LSCs. Indeed, dosing TUG1 downregulation by exposure to low and high CpG-TUG1-shRNAs (Fig. 7e *inset*) mimicked the dose-dependent differential triggering of *MIR300* anti-proliferative and pro-apoptotic activities exerted by different *CpG-miR-300* (Fig. 4); in fact, exposure of Ph^+^ cells to 100 nM *CpG-TUG1-shRNA* efficiently induced growth-arrest but modestly affected survival (Annexin V^+^ cells). Conversely, 500 nM *CpG-TUG1-shRNA* levels decreased CML-BC and CK-AML qLSC numbers in a *MIR300-*sensitive manner and further enhanced *CpG-miR-300*-induced apoptosis thereby causing a nearly complete loss of qLSCs and dividing progenitors (Fig. 7d, e). As expected, TUG1 overexpression (*TUG1 RNA*) antagonized *CpG-miR-300*-induced qLSC apoptosis in a *MIR300* dose-sensitive manner (Fig. 7e). Thus, the ability of CpG-anti-miR-300 to fully prevent TUG1-shRNA-induced apoptosis of CML/AML qLSCs (Fig. 7e) indicates that hypoxia-induced TGF*β*1-FoxM1-mediated signals increases TUG1 expression in CML/AML qLSCs to dynamically regulate quiescence by maintaining the amount of free functional *MIR300* at levels sufficient for arresting cell cycle but not for triggering PP2A-dependent apoptosis (Fig. 7a and Supplementary Fig. 7a).

While TUG1 seems to act identically in CML and AML qLSCs, the low and high TUG1 levels in CD34^+^ CML-BC and CK-AML progenitors, respectively (Fig. 7b), the inability of *CpG-anti-miR-300* to counteract TUG1-shRNAs-induced apoptosis of dividing CD34^+^ leukemic progenitors (Fig. 7e), and the nearly complete killing of CD34^+^ CML/AML progenitors by the combination of *CpG-miR-300* and *CpG-TUG1-shRNA* oligonucleotides (Fig. 8e) suggest that TUG1 activity in dividing leukemic blasts may involves requires the interaction with other miRNAs regulating leukemic cell proliferation/survival.

**Figure 8.**
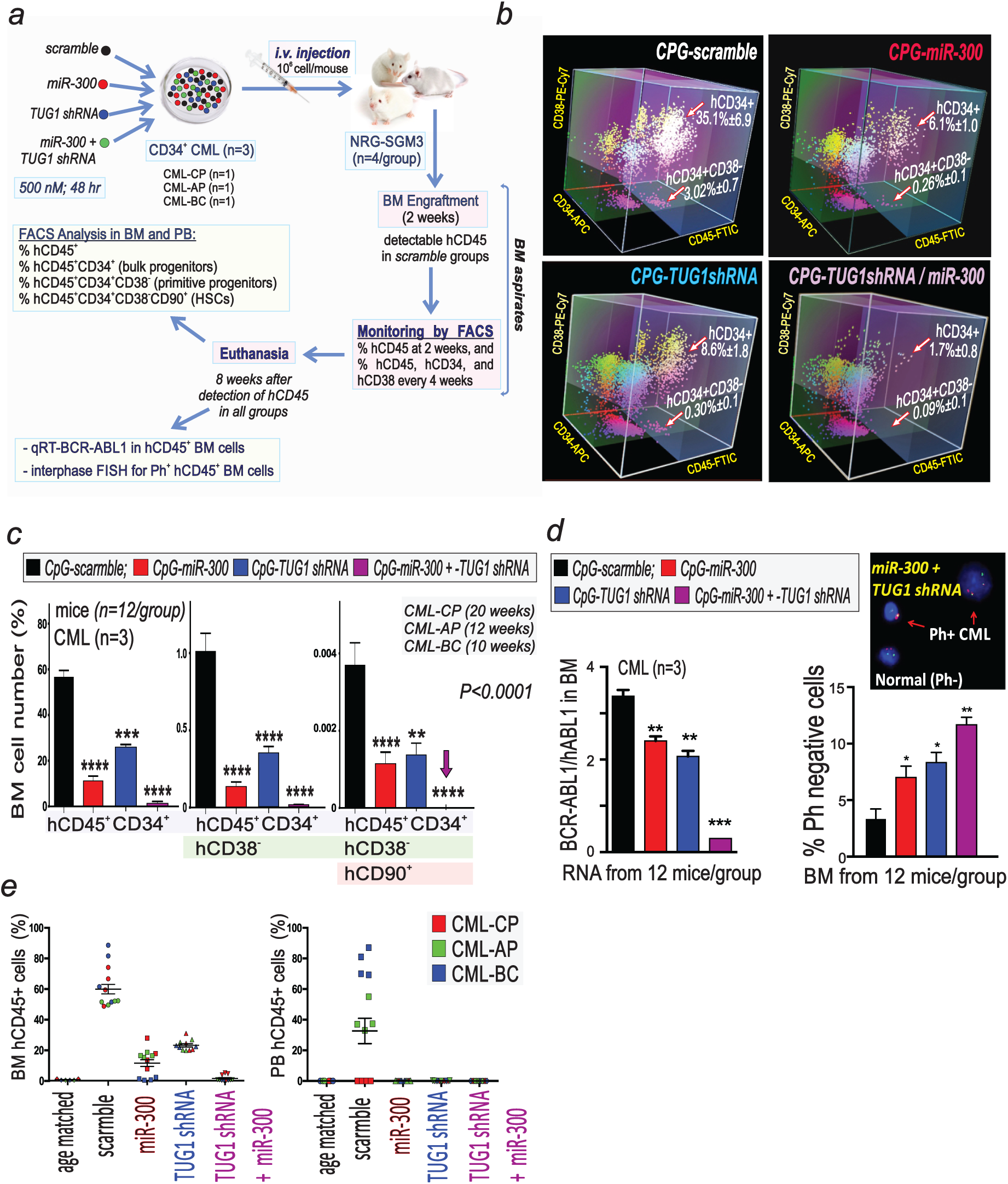
Halting *MIR300*-TUG1 interplay selectively eradicated quiescent LSCs in a PDX model of acute and chronic myeloid leukemias. (a) Xenotransplantation protocol of *ex vivo*-treated CD34^+^ CML cells in NRG-SGM3 mice (n=4 mice/treatment/patient sample). **(b, c)** Analysis of *CpG-MIR300-, CpG-TUG1shRNA-, CpG-TUG1shRNA+CpG-MIR300-* and *CpG*-*scramble-*treated CML cells from BM aspirates at 2-12 (3D-plots) and 10-20 weeks post-transplant quantitative analysis of CML cells stained with the indicated antibodies. **(d)** Evaluation at 10-20 weeks post-transplant of BCR-ABL1 transcripts by RT-qPCR (*left*) and of % Ph-negative (Ph^-^) cells by FISH (*right*) in total and FACS-sorted hCD45^+^BM cells, respectively. **(e)** Analyses of BM CML cells at 10-20 wk post-transplant. hCD45*^+^* cells (%) in BM (left) and PB (right) of mice transplanted with CML (CP, AP and BC) and treated with the indicated *CpG-ODNs*. Age-matched mice served as controls. Error bars indicate mean ± SEM.

Reportedly, TUG1 interacts with several miRNAs^45, 46, 51^ and bioinformatics analysis revealed that TUG1 may function as hub for tumor suppressor miRNAs with activities similar to that of *MIR300* (Supplementary Fig. 8c). By RNAseq, we found that 56 of these experimentally-validated TUG1-interacting miRNAs are differentially expressed in LSC-enriched CD34^+^CD38^-^ and committed CD34^+^CD38^+^ progenitor CML (CP and BC) compared to identical BM cell fractions from healthy (NBM) individuals (Supplementary Fig. 8a). Among these, 15 TUG1-interacting miRNAs are either aberrantly expressed or have a defined role in the regulation of proliferation and survival of AML/CML LSCs and/or progenitors (Supplementary Fig. 8a, c).

### Halting *MIR300*-TUG1 interplay selectively eradicated quiescent LSCs in a PDX model of acute and chronic myeloid leukemias

By using as previously described^52^ a PDX-based approach suitable for determining changes in leukemic LT-HSC survival *in vivo*, we assessed the effects of TUG1-dependent modulation of *MIR300* activity on BM-repopulating qLSCs and the therapeutic relevance of disrupting *MIR300*-TUG1 interplay. NRG-SGM3 mice (n=4/group) were transplanted with >95% Ph^+^ CD34^+^ CML-CP, -AP and -BC cells previously exposed for 48hr to 500nM *CpG-scramble* (control), *CpG-miR-300* and *CpG-TUG1shRNA* used as single agents or in combination (Fig. 8a). At 4 and 8 weeks after engraftment, BM hCD45^+^CD34^+^ progenitors and LSC-enriched hCD45^+^CD34^+^CD38^-^ cells were 75-97% reduced in all arms (Fig. 8b, c). Importantly, inhibiting and/or saturating TUG1 *MIR300*-sponging activity with *CpG-TUG1-shRNA* and *CpG-miR-300* resulted in ∼100% killing of leukemia-initiating (hCD45^+^CD34^+^CD38^-^CD90^+^) quiescent HSCs (Fig. 8c), barely detectable *BCR-ABL1* transcripts and 3.6-fold increased numbers of normal (Ph^-^) BM cells (Fig. 8d), and strongly reduced numbers of PB and BM CML (CP, AP and BC) hCD45^+^ cells (Fig. 8e). Notably, the lengthy CML-CP engraftment-time also indicated that decreased hCD34^+^ cell recovery resulted from reduced numbers of long-term BM-repopulating LSCs (LT-LSC) and not of transplanted CD34^+^ cells. Thus, disruption of *MIR300*-TUG1 balance suppresses acute and chronic myeloid leukemia development by selectively and efficiently eliminating nearly all drug-resistant leukemic but not normal quiescent HSCs and progenitors.

## DISCUSSION

Regardless of tumor type and stage, altered miRNA expression and PP2A tumor suppressor activity are tightly linked to tumorigenesis and impaired anti-cancer immunity^2, 5, 7^. Here we showed that therapeutically-exploitable post-transcriptional signals, which are initiated by the tumor-naïve and likely maintained by the tumor-reshaped BMM, controls CML and AML LSC entry/maintenance into quiescence and impairs NK cell immunity. This occurs through the induction of *MIR300*, a PP2A-activator and tumor suppressor miRNA with dose-dependent anti-proliferative and pro-apoptotic activities which are uncoupled and differentially regulated by the sponge activity of TUG1 lncRNA. Importantly, a dose-dependent target selection mechanism^53^ allows the sequential activation of *MIR300* anti-proliferative and pro-apoptotic functions through the inhibition of CDK6/CCND2 and SET, respectively, in qLSCs and leukemic progenitors.

### MIR300 role in LSCs and leukemic progenitors

Expression studies revealed that *MIR300* levels are downregulated in CML (CP and BC) and AML (CK) progenitors but not in qLSCs in which *MIR300* expression is strongly transcriptionally induced by hypoxia in a C/EBP*β*-dependent manner. Accordingly, Fanthom5 (http://fantom.gsc.riken.jp), TCGA (https://portal.gdc.cancer.gov) and tumor-specific miRNA sequence database^54, 55^ analyses (not shown) indicate that mature and/or precursor *MIR300* levels are higher in CD133^+^ CSCs but lower to undetectable in virtually all primary solid tumors and leukemia subtypes.

Restoration of *MIR300* expression arrested proliferation, expanded the G0/G1 cell cycle fraction, strongly impaired survival of dividing CD34^+^ leukemic stem/progenitor cells and was associated with downregulation of CDK6/CCND2, SET and other growth- and survival-promoting factors (e.g. Twist1 and CTNNB1). *MIR300*-induced downregulation of CCND1, Twist1 and CTNNB1 was also observed in several solid tumors; however, this was associated with growth arrest/apoptosis but, unclearly, also with the acquisition by tumor cells of a pluripotent stem cell phenotype^16, 20, 21, 46^.

Although *MIR300*-induced loss of CCND2/CDK6 (or CCND1/CDK4) likely represents the mechanism by which *MIR300* anti-proliferative activity contributes to stemness, since CCND2/CDK6 inhibition characterizes quiescent long-term HSCs (LT-HSCs) and is sufficient to arrest CD34^+^ leukemic progenitors in G0/G1^29, 30^, impaired expression of Twist1 and CTNNB1 unlikely determines either *MIR300* anti-proliferative and pro-apoptotic functions. By contrast it is conceivable that inhibition of SET accounts for *MIR300*-induced PP2A mediated apoptosis because i) SET downregulation is sufficient for triggering PP2A-mediated apoptosis of acute and chronic leukemia quiescent stem and progenitor cells^26^, and ii) expression of SET mRNAs lacking the high affinity *MIR300*-binding site efficiently rescued Ph^+^ cells from *MIR300*-induced PP2A-mediated apoptosis. Accordingly, bioinformatics analysis that integrates several algorithms and takes into account also miRNA levels in BM of normal and myeloid leukemic cells to identify miRNA targets^56–58^, indicated that inhibition of Twist1, CTNNB1 and other PP2A-regulated survival factors (Fig. 4a, Supplementary Fig. 2b, 3) will require levels of free *MIR300* higher than those suppressing SET. Thus, SET inhibition represents the main event leading to *MIR300*-induced PP2A activation.

Notably, the importance of CCND2/CDK6 and SET as key *MIR300* effectors is likely not limited to their *MIR300*-induced post-transcriptional downregulation (Supplementary Fig. 2a *right*). In fact, *SET*, *CCND2* and/or *CDK6* transcription, mRNA nuclear export, translation and/or protein stabilization/activation might also result from *MIR300*-induced inhibition of other PIP factors (e.g. hnRNPA1, JAK2) and their associated XPO1 nuclear export and SET-stabilizing SETBP1 proteins^26,28,59-61^.

Because qLSCs tolerate high *MIR300* levels and exhibit inactivation of PP2A and activation of JAK2, SET and CTNNB1^2, 23^, it is possible that *MIR300* pro-apoptotic activity is turned-off in qLSCs. This raises the questions of whether *MIR300* is required for LSC quiescence; how *MIR300* is regulated in qLSCs and leukemic progenitors; and, how qLSCs elude *MIR300*-induced apoptosis.

The requirement of *MIR300* for induction and maintenance of LSC quiescence is clearly demonstrated by i) the ability of anti-*MIR300* molecules to antagonize MSC-induced inhibition of leukemic cell proliferation, ii) impaired LTC-IC-driven colony formation in the absence of qLSC apoptosis in CD34^+^ cells exposed to *CpG-miR-300*concentrations that inhibit CCND2/CDK6 but not SET expression, and iii) inability of MSCs to induce SET downregulation in leukemic cells. Importantly, the notion that *MIR300*-dependent SET inhibition is induced by hypoxia in CD34^+^ leukemic stem/progenitor cells (Fig. 5b) and that SET, JAK2 and CTNNB1 are active in qLSCs^2, 23^, suggest that LSC entrance into quiescence is initiated prior to LSC niching into the BM endosteal area with the lowest O_2_ tension in which hypoxia-induced TUG1 inactivates *MIR300* pro-apoptotic function. Because exposure of leukemic CD34^+^ stem/progenitor cells to MSC CM and/or exosomes increases *MIR300* expression, suppresses their growth in a *MIR300*-dependent manner and double the number of CD34^+^ cells in G0 (Fig. 5c), it is likely that the transfer of MSC-derived exosomal *MIR300* into LSCs allows *MIR300*-induced CDK6/CCND2 downregulation and transition into quiescence.

Mechanistically, we showed that hypoxia induces *MIR300* transcription in MSCs and CD34^+^ leukemic cells through reduced LIP (inhibitory) and increased LAP1 (activatory) C/EBP*β* that binds/transactivates a hypoxia-sensitive regulatory element 109bp upstream the *MIR300* gene. Accordingly, C/EBP*β* was found to be expressed in LSCs, induced by hypoxia and to negatively regulate G1/S transition and SET expression^62, 63^.

To our knowledge, *MIR300* is the only cell context-independent tumor suppressor miRNA inducing LSC quiescence, inhibiting innate immunity and triggering LSC and leukemic progenitor apoptosis although other miRNAs regulating CSC self-renewal (e.g. miR-29, miR-125, miR-126, let-7) or survival of quiescent cells (e.g. mir-16 family) have been associated with CSC expansion and maintenance^4^. For example, miR-126, which targets CDK3, inhibits qLSC re-entry into cycle but does not induce quiescence and, importantly, it is not acting as a tumor suppressor in AML/CML LSCs^9, 10^. Less clear and controversial is the role of miR-221/222/223 in CSC quiescence since their expression is regulated by MSCs in a tumor- and cell type-specific manner but also increases in response to mitogenic stimuli to induce CDK4/CCND1-dependent cell cycle re-entry^4, 11^.

### MIR300 role in NK cells

*MIR300* is also the only tumor-naïve-induced tumor suppressor miRNA that inhibits NK cell-mediated innate anti-cancer immunity while promoting LSC quiescence. NK cells preferentially kill CSCs^64–66^, including BM-repopulating TKI-resistant BCR-ABL1^+^ qLSCs (Fig. 6b), and NK cell quantitative and functional impairment is a common event in detected in cancer at diagnosis^1, 15^. In CML, impaired NK cell immunity also associates with qLSC persistance in TKI-treated patients^24, 67^ whereas normal levels of activated NK cells characterize TKI-treated patients in sustained TFR^67^ and account for increased disease-free survival after T-cell-depleted stem cell transplant^64^, suggesting that NK cell-based therapies may lead to qLSC eradication.

Because *MIR300* levels are increased in circulating NK cells from CML patients at diagnosis (Fig. 6b) and, BM hypoxia- and MSC (CM and exosomes)-induced inhibition of NK cell proliferation and anti-tumor activity is essential for immunosurveillance^7^ and *MIR300* induction (Fig. 6c, d), it is plausible that NK cell inhibition in CML and, likely, in other cancer patients may arise from *MIR300*-mediated signals initiated by the naïve BMM ^7^ and overriding cytokine-driven NK cell activation ^35^. Mechanistically we showed that NK cell inhibition likely depends on BMM (hypoxia and MSCs)- induced C/EBP*β*-*MIR300* signals leading to CCND2/CDK6 and SET downregulation and that BMM-induced NK cell inhibition was recapitulated by *MIR300* mimics and rescued by *MIR300* RNAi (Fig. 6c-f).

Because loss of CCND2 and SET expression account for inhibition of NK cell proliferation^36^ and suppression of NK cell anti-tumor cytotoxicity^37^, it is conceivable that the tumor-reshaped BMM^68^ may exacerbate but not initiate *MIR300* or other signals that suppress cytokine-driven NK cell proliferation and cytotoxicity in leukemia patients. For example, the hypoxia-induced TGF*β*1 secretion by leukemic and other BM cells^7, 37, 69^ and the increase of TGFBR2 in NK cells ^69, 70^, may augment *MIR300*-induced CCND2 downregulation by uncoupling IL-2-dependent mitogenic and survival signals^71^. Seemingly, MSC exosome-induced pre-miR-155 (BIC) downregulation may contribute to *MIR300*-induced NK cell inhibition by antagonizing miR155-induced C/EBP*β* and PP2A downregulation leading to enhanced *MIR300* expression and further inhibition of MAPK/AKT-dependent NK cell inactivation, respectively^8, 35, 38, 39^. By contrast, TUG1 expression seems dispensable for human NK cell survival although, consistent with its activity as a *MIR300* sponge, it decreases in NK cells exposed to hypoxia and MSC-derived CM (Supplementary Fig. 6d, e).

Altogether the evidence that i) *MIR300* is critical for BMM-induced NK cell inactivation (Fig. 6), ii) clinically-relevant human NK-92 cells^34^ halve the number of TKI-resistant CML qLSCs, and that iii) ectopic miR-155 selectively and efficiently boost NK cell proliferation and cytotoxicity against mature tumor cells^35, 72^ and self-renewing LSCs (Fig. 6h), strongly supports the development of NK cell-based immunotherapies with agents that will prevent BMM inhibition and enhance NK cell-mediated selective qLSC and tumor cell killing by simultaneously boosting miR-155 and inhibiting *MIR300* activities.

### Biologic and therapeutic relevance of the MIR300-TUG1 interplay

TUG1 is an oncogenic lncRNA upregulated in CD34^+^ stem and progenitor cells from healthy individuals, CML and AML (Fig. 7 and Supplementary Fig. 7) and solid tumor patients in which it has strong diagnostic, prognostic and therapeutic relevance^45, 73^. In CML-BC and CK-AML qLSCs, TUG1 uncouples *MIR300* tumor suppressor function and in a dose-dependent manner selectively suppresses only *MIR300* pro-apoptotic activity. Experiments in which we used different *CpG-miR-300* or *CpG*-*TUG1-shRNA* doses (Fig. 4, 7) indicated that this unprecedented lncRNA function depends on the ability of TUG1 to keep the amount of free *MIR300* at levels sufficient for inducing growth arrest but not for triggering PP2A-mediated apoptosis. This implies that the nearly complete and selective (no effects on LT-HSC-driven normal hematopoiesis) killing of qLSCs and leukemic progenitors, which is induced *in vitro* and in PDXs by the combination of TUG1 shRNAs and *MIR300* mimetics (Fig. 7, 8), does not depend on loss of TUG1 survival signals but on the effect of the freed tumor suppressor miRNA (e.g. *MIR300*) on its mRNA targets. Accordingly, TUG1 loss was associated with G0/G1 arrest and apoptosis whereas its overexpression with induction of proliferation, EMT, drug-resistance and stemness in different solid tumors^46^, in which its activity was correlated with its sponging miRNA activity^46, 51^ and with the upregulation of mitogenic and survival factors (e.g. CCND1/2, CDK6, CTNNB1, Twist1) described as *MIR300* targets (Fig. 3). TUG1 activity seems to be *MIR300*-restricted in CML and AML qLSCs but not in leukemic progenitors in which TUG1-shRNAs induced apoptosis of anti-*MIR300*-treated CD34^+^ CML and AML cells (Fig. 7e). This is consistent with the association of TUG1 with dismal outcome in AML blasts^73^ and with the induction of TUG1 by AML1-ETO, MLL-AF4/9 and FLT3-ITD^73^ (Fig. 7e and Supplementary Fig. 7) and suggests that TUG1 may function as “*tumor suppressor miRNA warder*” controlling in a cell-type (LSCs and progenitors)-dependent the activity of specific subsets of tumor suppressor miRNAs (e.g. miRNAs inhibiting proliferation or leading to PP2A activation). This is likely facilitated by the recruitment of functionally-related miRNAs into specific miRNA-RBP ternary complexes (e.g. TUG1-hnRNPA1-SET/CCND2/CDK6)^59, 61, 74^. In this regard, we showed that 15 validated TUG1-interacting miRNAs are differentially expressed in CD34^+^CD38^-^ and CD34^+^CD38^+^ CML (CP and BC) BM cells and have defined roles in solid tumors and AML/CML stem/progenitor cells^75^ (Supplementary Fig. 8). Indeed, 96.4% and 44.3% of these miRNAs are predicted to shut down CML and AML oncogenic signals, and to share with *MIR300* the ability to regulate stemness, TGF*β* signaling and activate PP2A (Supplementary Fig. 8). Thus, it is conceivable that balanced TUG1-*MIR300* levels are essential for qLSC induction/maintenance whereas insufficient TUG1 expression will lead to qLSC and leukemic progenitor cells apoptosis by freeing miRNAs with similar tumor suppressor activities. Conversely, high TUG1 sponge activity will promote leukemic cell proliferation, survival and qLSC cell cycle re-entry (Supplementary Fig. 7a). However, an aberrant TUG1 increase at levels inhibiting also *MIR300* anti-proliferative activity in LSCs may induce LSC exhaustion by impairing entry into quiescence and forcing qLSC re-entry into cycle.

Mechanistically, we showed that TUG1 expression in qLSCs depends on hypoxia-induced TGF*β*1 secretion by CD34^+^ CML progenitors although we cannot exclude that hypoxia-induced TGFBR1/2 upregulation in LSCs and TGF*β*1 secretion by other BM cell types (e.g. MSCs)^48^ contributes to increase TUG1 levels to differentially regulate *MIR300* anti-proliferative and pro-apoptotic tumor suppressor functions.

TUG1 can also be induced by Notch signaling^76^ but the anti-TGF*β* Ab-mediated suppression of hypoxia-induced TUG1 levels in qLSCs argues against a Notch significant contribution to TUG1 induction. However, Notch signaling may contribute to qLSC expansion through the RBPJ-mediated miR-155 inhibition that may induce TUG1^50^ and *MIR300* expression by preventing FoxM1 and C/EBP*β* downregulation^77^, respectively. Indeed, we showed that TUG1 upregulation was dependent on hypoxia- and TGF*β*-induced FoxM1 expression in qLSCs. Accordingly, FoxM1 was described as an oncogene overexpressed in solid tumors and leukemias in which it promotes CSC quiescence, tumor cell survival and cell cycle progression^49, 78^ upon activation by ERK, CDK6 and PI3K-AKT signals (Supplementary Fig. 7a, d).

In conclusion, the tumor naïve BMM-induced *MIR300* tumor suppressor anti-proliferative and PP2A-activating functions support leukemogenesis through the induction of LSC quiescence and inhibition of qLSC killing by cytokine-activated NK cells, respectively. This may represent the initial step leading to formation and initial expansion of the qLSC pool. Once established, the leukemic clone will reshape the BMM to further support disease development and progression. TUG1-*MIR300* interaction play a central role in this process levels since altering its ratio leads to the nearly complete and selective PP2A-dependent eradication of acute and chronic myeloid leukemias at qLSC levels *in vitro* and in PDXs. Thus, this work not only highlights the therapeutic importance of altering *MIR300* levels in anti-LSC and NK cell-based approaches for leukemias and, likely, several solid tumors with similar *MIR300* and TUG1 expression patterns but also indicates that the activity of a tumor suppressor can be exploited by cancer stem cells to preserve their ability to induce and maintain cancer.

## METHODS

### Cell lines and primary cells

*Cell lines:* Ph^+^ CML-BC K562 and LAMA-84, BM MSC-derived HS-5^79^ and the clinically-relevant NK-92^34^ cells were cultured in RPMI-1640 medium. NK-92 cultures were supplemented with 150 IU/ml rhIL-2 (Hoffman-La Roche Inc.). The amphotropic-packaging 293T and Phoenix cells were cultured in Dulbecco modified Eagle medium (DMEM). All tissue culture media were supplemented with 10-20% heat-inactivated FBS (Gemini; Invitrogen), 2 mM L-glutamine and 100 U/ml Penicillin/StreptoMycin (Invitrogen). The 32D-BCR-ABL and the IL-3-dependent parental 32Dcl3 mouse myeloid precursors were cultured in Iscove modified Dulbecco Medium (IMDM) supplemented with 10% FBS and rmIL-3 (2ng/ml, Peprotech). *Primary cells:* Human hematopoietic progenitor (CD34^+^) and stem cell-enriched fractions (CD34^+^ CD38^-^) from healthy and leukemic individuals were isolated from bone marrow (BM), peripheral (PB) or umbilical cord (UCB) blood. Prior to their use, cells were kept (18h) in StemSpan^TM^ CC100 cytokines-supplemented SFMII serum-free medium (StemCell Technologies). Human BM MSCs (hMSCs) from healthy individuals were isolated from BM cells by Ficoll-Hypaque density gradient centrifugation followed by culture in complete human MesenCult^TM^ proliferation medium. MSC purity (>99%) was determined by flow cytometry. Human CD56^+^CD3^-^ NK cells (purity >95%) from healthy (UCB or PB) and CML individuals were FACS and/or magnetic (Miltenyi Biotech Inc.) cell sorted, or RosetteSep Ab-purified (StemCell Technologies) as described^37^. Frozen leukemia (CML and AML) specimens were from the Leukemia Tissue Banks located at The University of Maryland (UMB, Baltimore, MD); The Ohio State University (Columbus, OH), Maisonneuve-Rosemont Research Centre, Montreal (Quebec, Canada); Hammersmith Hospital, Imperial College (London, UK); Policlinico Vittorio Emanuele (Catania, Italy); University of Utah, (Salt Lake City, UT); Hematology Institute Charles University (Prague, Czech Republic) and Aarhus University Hospital (Aarhus, Denmark); fresh UCB, NBM and PB (CML and healthy individuals) samples were purchased (Lonza Inc.) or obtained from UMB hospital and Maisonneuve-Rosemont Research Centre, Montreal (Canada).

Cells were treated for the indicated time and schedule with 1-2 μM imatinib mesylate (IM; Novartis), 5 μM 5-Aza-2′-deoxycytidine (5-Aza; Sigma), 250-500 nM *CpG-scramble*, *-MIR300*, - *anti-MIR300* and -*TUG1shRNA* oligonucleotides (ODNs) (Beckman Research Institute, City of Hope), 1.25 μg/ml anti-TGF*β* neutralizing antibody (1D11; R&D Systems), 10 ng/ml rhIL-12 and 100 ng/ml rhIL-18 (R&D Systems). Where indicated, cells were cultured for the indicated times in hypoxic conditions (1% O_2_), HS-5 and hMSC CM (100% vol/vol), or in medium supplemented with MSC-derived exosomes (50-100 μg/ml). During treatments, viable cells were enumerated by Trypan Blue exclusion test.

Mouse NK1.1^+^CD3^-^ NK cells and Lin^-^Sca^+^Kit^+^ (LSK) cells were purified from spleen of 8-10wk-old healthy and leukemic (6-8 wk-induced) SCL-tTA-BCR-ABL tg mice^80^ and wt and *lck*-miR-155-tg mice^39^ by microbeads negative selection (Miltenyi Biotech Inc.) and cell sorting.

### Plasmids

<u>pCDH-*MIR300* (hsa-*MIR300*) and pCDH-miR-381 (hsa-miR-381-3p)</u> pCDH-*MIR300* was generated by subcloning a 490bp NheI-BamHI PCR fragment encompassing the mature hsa-*MIR300* into the pCDH-CMV-MCS-EF1-copGFP-puro (SBI) vector. Sequence was confirmed by sequencing. pCDH-miR-381 was generated by subcloning a double-stranded (ds) synthetic ODN containing the human pre-miR-381 sequence flanked by *NheI* and BamHI restriction sites at the 5’- and 3’-end, respectively, into the pCDH-CMV-MCS-EF1-copGFP-puro vector. <u>pSIH-H1-Zip-</u> *<u>MIR300</u>*: To knockdown *MIR300*, a 5’-EcoRI and 3’-BamHI-flanked *MIR300* antisense dsODN was directionally cloned into the pSIH1-H1-copGFP vector (SBI). <u>pSIH-H1-copGFP-shTUG1</u> <u>(TUG1 shRNA)</u>: a shRNA cassette containing the targeted nt 4571 to 4589 of hTUG1 RNA^76^ was subcloned into pSIH-H1-copGFP vector (SBI). A non-functional scrambled TUG1 shRNA was used as a control. <u>pLenti-TUG1:</u> the hTUG1 into the pLenti-GIII-CMV-GFP-2A-Puro-based vector was from Applied Biological Materials Inc. <u>pGFP/Luc-based *MIR300* promoter constructs:</u> p513-GFP/Luc, p245-GFP/Luc, and p109-GFP/Luc were generated by cloning the -522 bp, -245 bp and -109 bp regions upstream the hsa-*MIR300* gene locus (Sequence ID: <u>NC_000014.9</u>) into pGreenFire1 (pGFP/Luc, pGF1; SBI) reporter vector. The regions upstream the hsa-*MIR300* gene were PCR amplified from K562 DNA and cloned into the pGF1 EcoRI-blunted site. p109-C/EBPBmut-GFP/Luc: an EcoRI-containing double-stranded synthetic oligonucleotide spanning the -109bp *MIR300* regulatory (region and containing the T/G and C/G mutated C/EBPβ consensus binding sites located at positions -64 and -46, respectively was subcloned into EcoRI-digested pGF1 vector. <u>pCDH-Flag-SET</u>, fluorescent ubiquitination-based cell cycle indicator reporter <u>pCDH-FUCCI2BL</u>, MigR1-ΔuORF-C/EBP*β*-ER^TAM^ and <u>MigR1-ΔuORF-C/EBP</u>*<u>α</u>*<u>-HA</u> constructs were described^32, 81–83^. The pCDH-Flag-SET 3^’^UTRwt-GFP and <u>pCDH-Flag-SET</u> Δ3^’^UTR-GFP: wild-type (627bp) or 3**’**-deleted (513bp) SET 3**’**UTRs were PCR amplified from K562 cDNA using a common *EcoRI-linked* 5’-primer and specific BamHI-linked 3’-primers for the wild-type and deleted *SET* mRNA 3’UTR. The PCR products were cloned into the pCDH-Flag-SET plasmid and sequenced.

### MSC Conditioned Medium (CM) and Exosomes purification

HS-5- and hMSC-derived CM was obtained by culturing cells in complete RPMI (24h) and StemSpan^TM^ SFMII (48h) medium, respectively. Alix^+^CD63^+^ exosomes were purified by differential centrifugation (3,000xg, 15 min; 10,000xg, 10 min; and 100,000xg, 70 min) from HS-5 CM cultured in exosome-free FBS-supplemented medium^84, 85^. When required, exosomes were precipitated (Exo-Quick; SBI) prior to isolation by ultracentrifugation.

### Flow cytometry and cell sorting

CD34^+^ and CD34^+^CD38^-^ fractions were magnetic-(CD34 MicroBead kit; Miltenyi Biotec) and/or FACS-(*α*CD34 APC/PE and *α*CD38 PE/Cy7 Abs, BD Biosciences) purified. Primary human NK cells were sorted using Alexa fluor 488 *α*hCD56, PE-Cy5 *α*hCD19 and PE-Cy7 *α*hCD3 Abs (BD Bioscience) and their purity assessed by PE *α*CD56 (Beckman Coulter, Life Sciences) and APC-eFluor780 *α*CD3 (eBioscience) Ab staining. Mouse NK cells were sorted using *α*NK1.1 APC and *α*CD3 FITC Abs (BD Bioscience) and LSK cells using *α*Lin^-^ (lineage cocktail) Pacific Blue (Biolegend), *α*c-Kit PE-Cy7 and *α*Sca1 APC Abs (eBioscience). hMSCs purity was assessed by flow cytometry using anti-CD34 and CD45 FITC, CD73 PE-Cy7, CD105 Alexa 647, CD44 Percp-Cy5.5 and CD90 PE Abs (BD Biosciences). Apoptosis was quantified by FACS upon staining cells with PE Annexin-V and 7-ADD (BD Biosciences). Cells were sorted using the FACS Aria II (BD Biosciences). Data acquisition and analyses were performed at the UMBCCC Flow Cytometry Facility. Data from LSRII or CANTO II flow cytometers (BD Biosciences) were analyzed by using either the FlowJo v8.8.7 or Diva v6.1.2 software.

### Lentiviral and retroviral transduction

Lentiviral or retroviral particles were produced by transient calcium phosphate transfection (ProFection mammalian transfection System; Promega) of 293T and Phoenix cells, respectively^2^. Briefly, viral supernatant was collected 24 and 48h after transfection and directly used or concentrated by PEG precipitation. Viral titer was determined by flow cytometry by calculating number of GFP^+^ 293T cells exposed (48h) to different viral dilutions. One-to-three spinoculation rounds were used for cell line infection whereas a single spinoculation with diluted viral supernatants (MOI=6) was used for primary cells. Viral supernatants were supplemented with polybrene (4 μg/ml). FACS sorting or puromycin selection was initiated 48h after transduction.

### Cell cycle analysis

Cell cycle analysis was performed by 4^’^,6^’^-diamidine-2-phenylindole (DAPI)/Ki-67 staining of *CpG-scramble-* or *CpG-MIR300*-treated (500nM; 72h) primary CD34^+^ CML (CP and BC) and UBC cells as described^83^. Briefly, cells were fixed, permeabilized (Cytofix/Cytoperm kit, BD Biosciences) and, Ki-67 PE (Biolegend) and DAPI (Sigma) stained. Alternatively, cell cycle analysis was performed on pCDH-Fucci2BL-infected cells and subjected to flow-cytometric analysis. LAMA-FUCCI2BL^+^ CML-BC cells were exposed to 500nM *CpG-scramble* and *CpG-miR-300*oligonucleotides after being G1/S synchronized (4-6 µM aphidicolin, 6h). *CpG-oligo*-treated (48h) FUCCI2BL^+^ cells were subjected to live cell imaging for 48h and analyzed as described^83^. Briefly, *CpG-scramble-* or *CpG-MIR300*-treated-treated cells in G0/G1 were mVenus^-^ PE-Texas-Red^+^, G1/S cells were mVenus^+^ PE-Texas-Red^+^, and S/G2/M cells were mVenus^+^ PE-Texas-Red^-^.

### Long-term culture-initiating cell (LTC-IC) and colony-forming cell (CFC)/replating assays

CD34^+^ CML (CP and BC), AML and UCB cells were lentivirally-transduced/GFP-sorted and/or treated (500 nM, 3d) with *CpG-ODN* in order to ectopically express either *MIR300* or anti-*MIR300* RNAs prior to use them in LTC-IC and CFC assays^2^. Vector-transduced and CpG-scramble-treated cells served as controls. *LTC-ICs:* 2×10^5^ CML and 2×10^3^ UCB CD34^+^ cells were cultured with a 1:1 mixture of irradiated (80 Gy) IL-3/G-CSF–producing M2-10B4 and IL-3/KL-producing SI/SI murine fibroblasts in MyeloCult H5100 (StemCell Technologies) supplemented with hydrocortisone. Medium was replaced after 7 days, followed by weekly half-medium changes, and fresh 500 nM *CpG*-*ODN* where required. After 6 wk, adherent and floating cells were harvested, and 5×10^3^ CML and 2×10^3^ for UCB cells were plated into cytokine (KL, G-CSF, GM-CSF, IL-3, IL-6)-supplemented MethoCult H4435. LTC-IC-derived colonies were scored after 14 days. *CFCs and CFC/Replating assays*: CD34^+^ and CD34^+^ CD38^-^ CML/AML (5×10^3^) and UCB (2×10^3^) cells were seeded in MethoCult H4435 supplemented with KL, G-CSF, GM-CSF, IL-3, IL-6. After 2 wk, colonies were scored and, if necessary, 10^4^ cells underwent two rounds of serial replating/scoring.

### CFSE (or eFluor670)-mediated tracking of cell division

Carboxyfluorescein diacetate succinimidyl diester (CFSE; CellTrace CFSE Proliferation Kit; Invitrogen) or eFluor670 (Cell proliferation Dye eFluor670; eBioscience)-stained cells were FACS-sorted to isolate the highest fluorescent peak, treated as indicated, harvested after 4-5 days and counterstained with Near-IR fluorescent Dye (Live/Dead Cell Stain Kit, Invitrogen) to determine the number of viable dividing (CFSE/NearIR^-^) and quiescent cells (CFSE^max^/NearIR^-^). Where indicated, cells were stained with anti-CD34 APC and sorted into dividing and quiescent sub-populations. Quiescent (CFSE^max^CD34^+^) and dividing cells in each peak were reported as a fraction of the initial number of CD34^+^ cells^2^. For NK cells, 4 and 7 day-cultured CFSE-labeled NK-92 and primary NK cells, respectively, were *α*CD56 APC Ab- and Aqua Fluorescent Dye (Live/Dead Cell Stain Kit, Invitrogen)-stained, and proliferation quantitated as fold changes of CFSE Mean Fluorescence Intensity (MFI) in living CD56^+^ NK cells.

### Live Cell Imaging and Confocal Microscopy

Wide field fluorescence imaging of *CpG-ODN*-treated pCDH-FUCCI2BL-transduced LAMA-84 cells was executed with an Olympus Vivaview digital camera, maintained at humidified atmosphere 37°C with 5% CO_2_, mounted into a microscope-equipped incubator with green and red fluorescence filters. Emission spectral ranges were green-narrow (490-540 nm) and red-narrow (570-620 nm). Imaging was carried out for 24 or 48 hours. Confocal fluorescence images were acquired using a LEISS LSM510 inverted confocal microscope equipped with 40X/1.2-NA water immersion objective with 0.55micron pixel size. Cell cycle phases were determined by plotting average red and green fluorescent intensity over time using Prism 6.0d software. The Vivaview raw images were reconstructed using ImageJ v5.2.5 and Adobe CS6 software.

### PP2A phosphatase assay

PP2A assays were performed using the malachite green–based PP2A immunoprecipitation (IP) phosphatase assay kit (Millipore) as described^2^. Briefly, protein lysates (50μg) in 100 μl of 20mM HEPES pH 7.0/100 mM NaCl, 5 μg *α*PP2Ac Ab (Millipore) and 25 μl protein A–agarose was added to 400 μl of 50 mM Tris pH 7.0, 100 mM CaCl_2_. Immunoprecipitates were carried out at 4°C for 2h and used in the phosphatase reaction according to the manufacturer’s protocol.

### Luciferase Assay

Transcriptional activity of the intergenic 513 bp region upstream the *MIR300* gene was investigated by luciferase assay using the pGreenFire-Lenti-Reporter system (pGF1; SBI) in cell lines and primary CD34^+^ CML-BC cells. Briefly, cells were transduced with constructs containing wild type full-length 513 (p513-GFP/Luc) and, deleted 245 (p245-GFP/Luc) and 109 (p109-GFP/Luc) base pair-long region upstream the *MIR300* gene locus, or with the p109-GFP/Luc construct mutated in the two C/EBPβ binding sites (p109-CEBP*β*mut-GFP/Luc). The empty vector (pGFP-Luc; pGF1) was used as a negative control. After 48h, cells were lysed and luciferase activity was determined by Pierce Firefly Luciferase Flash Assay Kit (Thermo Scientific).

### Cytotoxicity assays

A flow cytometry-based killing assay was performed using K562 cells as targets^86^. Briefly, HS-5-derived CM (100% vol/vol)- or exosomes (50-100 ug/ml)-preconditioned (36h in 50-150 IU/ml IL-2) NK-92 cells were co-incubated (3.5h) with CFSE-labeled K562 cells at ratio 5:1. Killing was evaluated by assessing the percentage of Annexin-V^+^ cells on CD56^-^ and/or CFSE^+^ target cells. When *CpG-ODNs* were used, NK-92 cells were preconditioned (36h) in medium lacking IL-2. NK-92 cell-killing of qLSCs was performed using CFSE-labeled CD34^+^ CML-BC cells cultured (4d) in cytokine-supplemented StemSpan^TM^ II serum-free medium exposed (18h) to NK-92 cells. Spontaneous cytotoxicity toward qLSCs was determined by FACS-mediated evaluation of LSC numbers in Annexin V^neg^CD34^+^CFSE^max^ cells before and after exposure to NK-92 cells. wt and miR-155-tg murine NK cell-mediated killing of LSK cells was determined by assessing CFC/replating efficiency in FACS-sorted CFSE^+^ Lin^-^Sca1^+^Kit1^+^LSK cells exposed to NK1.1^+^CD3^-^NK cells (18h).

### Real time RT-PCR

Total RNA was extracted using miRNeasy micro Kit (Qiagen Inc.) or Trizol (Invitrogen) and reverse transcribed using a standard cDNA synthesis or the TaqMan MicroRNA Reverse Transcription Kit with mouse and/or human-specific sets of primers/probe for *BCR-ABL1*, *CEBPB, FOXM1*, *TUG1*, *IFNγ, pri-MIR300, MIR300, miR-381, pre-miR-155 (BIC), 18S, RNU44, RNU6B* and *snoRNA202* (Applied Biosystems). PCR Reactions were performed using a StepOnePlus Real Time PCR System (Applied Biosystems). Data were analyzed according to the comparative C_T_ method using *RNU44, RNU6B, 18S* and *snoRNA202* transcripts as an internal control. Results are expressed as fold change of mean±SEM.

### Immunoblot analysis

Lysate from cell lines and primary cells were subjected to SDS-PAGE and Western blotting as described^2^. The antibodies used were: anti-Actin, anti-Myc, anti-C/EBPβ, anti-Alix, anti-Twist, and anti-CD63 (Santa Cruz Biotechnology); anti-GRB2 (Transduction Laboratories); anti-JAK2, anti-β-catenin, anti-FoxM1, anti-CDK6, anti–pJAK2^Y1007/1008^ and anti-CCND2 (Cell Signaling); anti-SET (Globozymes); anti-ABL (Ab-3), anti-CCND1/2, anti-PY (4G10) and anti-PP2Ac^Y307^ (EMD); anti-hnRNPA1 (Abcam); and anti-Flag (M2; Sigma).

### LSC engraftment and disease development in NRG-SGM3 mice

*In vivo* analysis of *ex vivo*-treated BM-repopulating LSCs was performed as described^52^. 10^6^ BM CD34^+^ CML (n=3; >95% Ph^+^; CP, AP and BC) cells, treated (500 nM, 48h) *ex vivo* with *CpG*-ODNs, were intravenously (iv) injected (4 mice/treatment group/patient sample) into sub-lethally irradiated (2.6Gy) 6-8 week-old NOD.Cg-*Rag1^tm1Mom^ Il2rg^tm1Wjl^* Tg(CMV-IL3, CSF2,KITLG)1Eav/J (NRG-SGM3; Jackson Laboratory). Engraftment was assessed by anti-human CD45 (BD Biosciences) flow staining of intra-femur BM aspirates^2^ at 2 wk in *CpG-scramble* control groups and at 2, 4 and 12 wk post-transplant in *CpG-MIR300*, -*TUG1shRNA* and *MIR300+TUG1shRNA* CML-BC, AP and CP, respectively. Disease evolution and effect of the various *CpG-ODNs* were assessed by FACS-mediated analysis of hCD45^+^ cells in BM and PB, and of hCD45^+^CD34^+^ proliferating progenitors, hCD45^+^CD34^+^CD38^−^ and hCD45^+^CD34^+^CD38^−^CD90^+^ CSC-enriched cell fractions with repopulating activity in BM aspirates at engraftment time and/or 8 wk post-engraftment (end point). BCR-ABL1 transcript levels were monitored by RT-qPCR-mediated (Ipsogen) analysis of *BCR-ABL1*/human *Abl1* ratios in total RNA samples derived from total BM of mice euthanized at 8 wk post-engraftment. Interphase FISH was performed on hCD45^+^-sorted BM cells, obtained at 8 wk post-engraftment, by denaturing (75°C, 2 min) slides in 70% formamide/2x standard sodium citrate (SSC) and ET-OH dehydrated. Interphase cells were hybridized (18h, 37°C) with BCR-ABL1 dual-color double-fusion probe set (10 UL; Cytocell) and DAPI counterstained. Slides were analyzed using an Olympus fluorescence microscope, and images taken with a CCD camera using a FISH imaging software (MetaSystems). 200-interphase cells/sample were analyzed and FISH negativity (Ph^-^) was defined as the absence of BCR-ABL fusion signal.

### Oligonucleotides and Primers

The partially-phosphothioated (*) 2′-O-Methyl ODNs were linked using 5 units of C3 carbon chain linker, (CH2)_3_ (X). The ODNs were synthesized by the DNA/RNA Synthesis Laboratory, Beckman Research Institute of the City of Hope.

#### CpG-oligonucleotides and probes

*CpG-MIR300*: 5^’^-g*g*tgcatcgatgcagg*g*g*g*gxxxxxuauacaagggcagacucucucu-3^’^

*CpG-anti-MIR300*: 5^’^-g*g*tgcatcgatgcagg*g*g*g*gxxxxxagagagagucugcccuuguaua-3’

CpG-TUG1 shRNA: 5^’^-g*g*tgcatcgatgcagg*g*g*g*gxxxxxuuacucugggcuucugcac-3’

*CpG-scramble*: 5^’^-g*g*tgcatcgatgcagg*g*g*g*gxxxxxguagaaccguacucgucacuua-3^’^

#### pCDH-CMV-MCS-EF1-copGFP-puro miRNA constructs

*MIR300(F):* 5’-gctagctgtgactagttgtaccttag-3’; *MIR300 (R):* 5’-gatcctctcttccagaaagttcttg-3’

*miR-381(+):*5’-ctagctacgcaaagcgaggttgccctttgtatattcggtttattgacatggaatatacaagggcaagctctctgtgagtag-3’

*miR-381(-):*5’-gatcctactcacagagagcttgcccttgtatattccatgtcaataaaccgaatatacaaagggcaacctcgctttaagta-3’

#### pSIH1-H1-copGFP constructs

Zip-*MIR300* ODN: 5’-gatatgttcccgtctgagagagagaaggacagtcttctctctcagacgggaacatataaaaacttaa-3’

shTUG1(+): 5’-gatccgtgcagaagcccagagtaattcaagagattactctgggcttctgcactttttg-3’

shTUG1 (-): 5’-aattcaaaaagtgcagaagcccagagtaatctcttgaattactctgggcttctgcacg-3’

#### pCDH-Flag-SET constructs

SET3^’^UTRwt-GFP(F): 5’-gaattctagcttttttcctcctttctctgtata-3’;

SET3^’^UTRwt-GFP(R): 5’-cggatccgtatacaagtcaaact-3’

SETΔ3^’^UTR-GFP(F): 5’-ggatccagagaaaagcatccaa-3’;

SETΔ3^’^UTR-GFP(R): 5’-cggatccgtatacaagtcaaact-3’

#### pGreenFire1

p522-pGF1(F): 5’-ccggaattccggcacgttttcagtatcaaatgct-3’

p245-pGF1(F): 5’-ccggaattccggtgaacctcttttactgtgactagttg-3’

p109-pGF1(F): 5’-ccggaattccggggtgtgctgctctcaccat-3’

Common(R): 5’-tgctctagagcaaatgatggcagtgacaggaa-3’

#### p109-C/EBPβ mutant construct

(+) strand ODN: 5’-aattc ggtgtgctgc tctcaccatg cagatcccat ctgtgtctct aaggc**t<u>g</u>gc**t cct**g<u>g</u>ag**ctg gtgggaactt agtcacagag gaaatggcct tcctgtcact gccatcattg-3’

(-) strand ODN: 5’-caatgatggc agtgacagga aggccatttc ctctgtgact aagttcccac cagctccagg agccagcctt agagacaca gatgggatct gcatggtgag agcagcacac cgaatt-3’

### Bioinformatics tools

Statistically significant (p<0.05 with FDR correction) predicted and validated hsa-*MIR300* mRNA targets according to mRNA target function and *MIR300* doses were identified using DIANAmicroT-CDS (diana.imis.athena-innovation.gr), ComiR (benoslab.pitt.edu/comir), (cosbi.ee.ncku.edu.tw/CSmiRTar/) and mirDIP 4.1 (ophid.utoronto.ca/mirDIP/). Kyoto Encyclopedia of Genes and Genomes (KEGG) and Gene Ontology (GO) analyses to define the biological pathways and the functional roles of miRNA-300 and TUG-1 targets were performed using miR-Path v.3 (snf-515788.vm.okeanos.grnet.gr). miRTar algorithm (miRTar.mbc.nctu.edu.tw/) was used to predict mRNA targets of wild type and ADAR-1-edited miR-381-3p. *MIR300* and TUG-1 individual gene expression profiles were obtained from curated datasets in the Gene Expression Omnibus (GEO) repository (ncbi.nlm.nih.gov/geoprofiles/). TUG1 levels in normal myelopoiesis and different AML subtypes were analyzed from BloodSpot database (servers.binf.ku.dk/bloodspot/) of healthy and malignant hematopoiesis. Integration of StarBase v2.0 (starbase.sysu.edu.cn/starbase2/index.php) database with the RNAseq MiRbase data from CD34^+^CD38^-^, CD34^+^D38^+^ CML and NBM cells was used to identify TUG-1 sponged miRNAs.

### Statistics

*P* values were calculated by Student’s t-test (GraphPad Prism v6.0). Results are shown as mean ± SEM. A *P* value less than 0.05 was considered significant (∗*P*< 0.05, ∗∗*P* < 0.01, ∗∗∗ *P* < 0.001, ∗∗∗∗ *P* < 0.0001). Mixed models’ approach, the split-plot design, was used to assess differences in percent cell across three treatment groups. The percent cell change was estimated using a model with two main effects for treatment and stage of a cell cycle, as well as their interaction. Tests of fixed effects revealed that interaction of treatment and stage of a cell cycle is highly significant, p=0.01, the two main effects, treatment and stage, should be interpreted with caution, the p-values are 1.0 and <0.0001, respectively. The differences in average percent cell were tested across groups and sliced at a particular stage of cell cycle.

### Study approval

Patient samples were banked at different Institutions and not collected for this study, which was carried out with a waiver of informed consent and approval from University of Maryland IRB. UCB units were collected at University of Maryland Medical Center with IRB approved protocol and written patient consent. Animal studies were performed according to IRB- and IACUC-approved protocols.

## ACKNOWLEDGEMENTS

We thank Goloubeva O. for statistical data analysis; Underwood K.F., Mauban J.R.H., Pomicter T., Duong V.H., Emadi A., Eiring A.M, and Glynn-Cunningham N.M. for patient sample procurement, technical assistance, reagents, and/or critical reading of the manuscript.

## AUTHORS CONTRIBUTIONS

Conceptualization and Funding Acquisition: D.P. Writing Original Draft: D.P. Investigation (>75% effort): G.S., R.T. (equal contribution) Supporting Investigation (<25% effort): L.S., S.W., J.J.E., J.H., P.N., E.A.K. K.S., M.M.M., Y.Z.; Provided Resources: C.G.P., G.P., C.H.M.J., F.S., P.V., G.N., P.M., A.R., R.G., D.C.R., M.G., P.H., M.D., G.F., C.H., F.D., D.M., J.A., G.M., J.Q., K.M.P., Y.Z., X.F., M.R.B. and B.C.

## SUPPLEMENTARY FIGURES

**Supplementary Figure 1.**
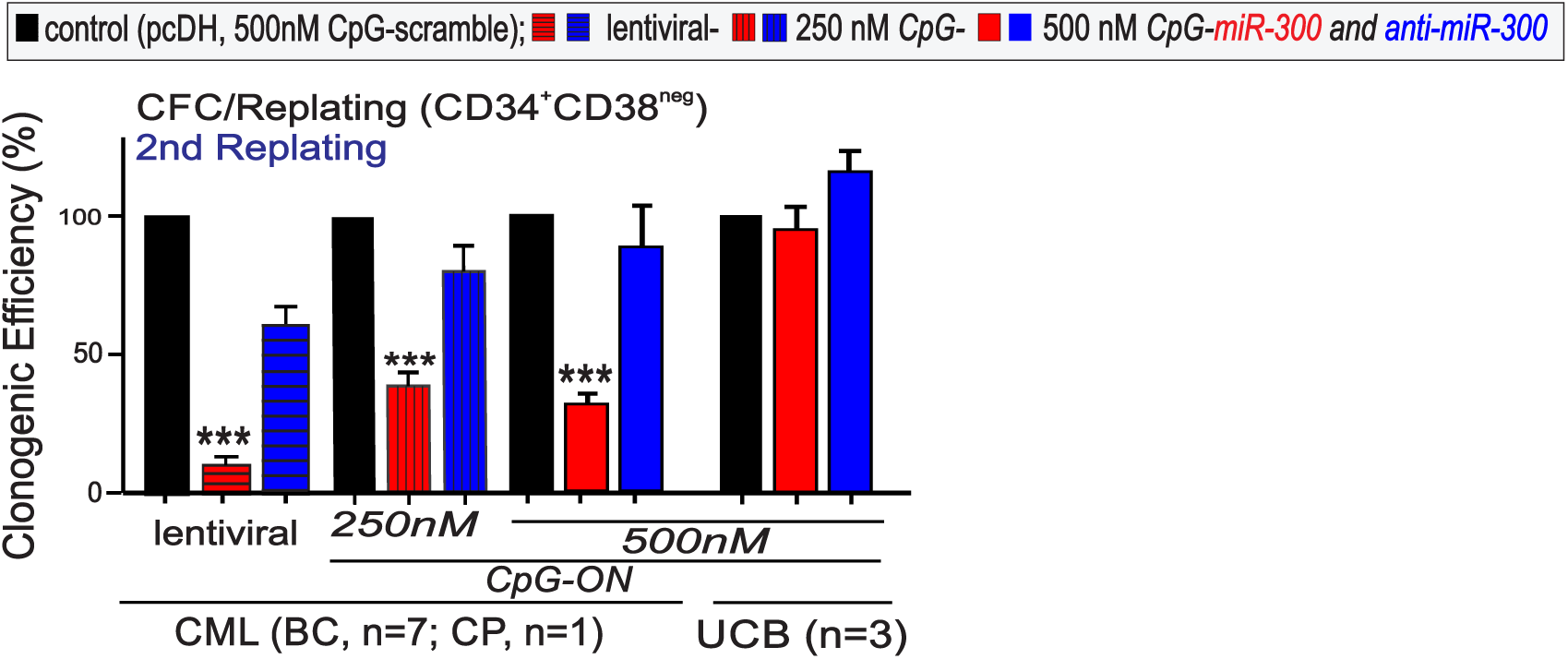
MIR300 activity in quiescent leukemic stem and progenitor cells. CFC-replating assays shows effects of lentiviral-mediated ectopic *MIR300* expression, 250 nM and 500 nM *CpG-miR-300*on serial replating activity (2^nd^ replating) of leukemic chronic and acute CML and normal UCB CD34^+^CD38^-^ HSC-enriched cell fractions. Infection with lentiviral empty vector and treatment with CpG-anti-*MIR300* and CpG-scramble served as controls.

**Supplementary Figure 2.**
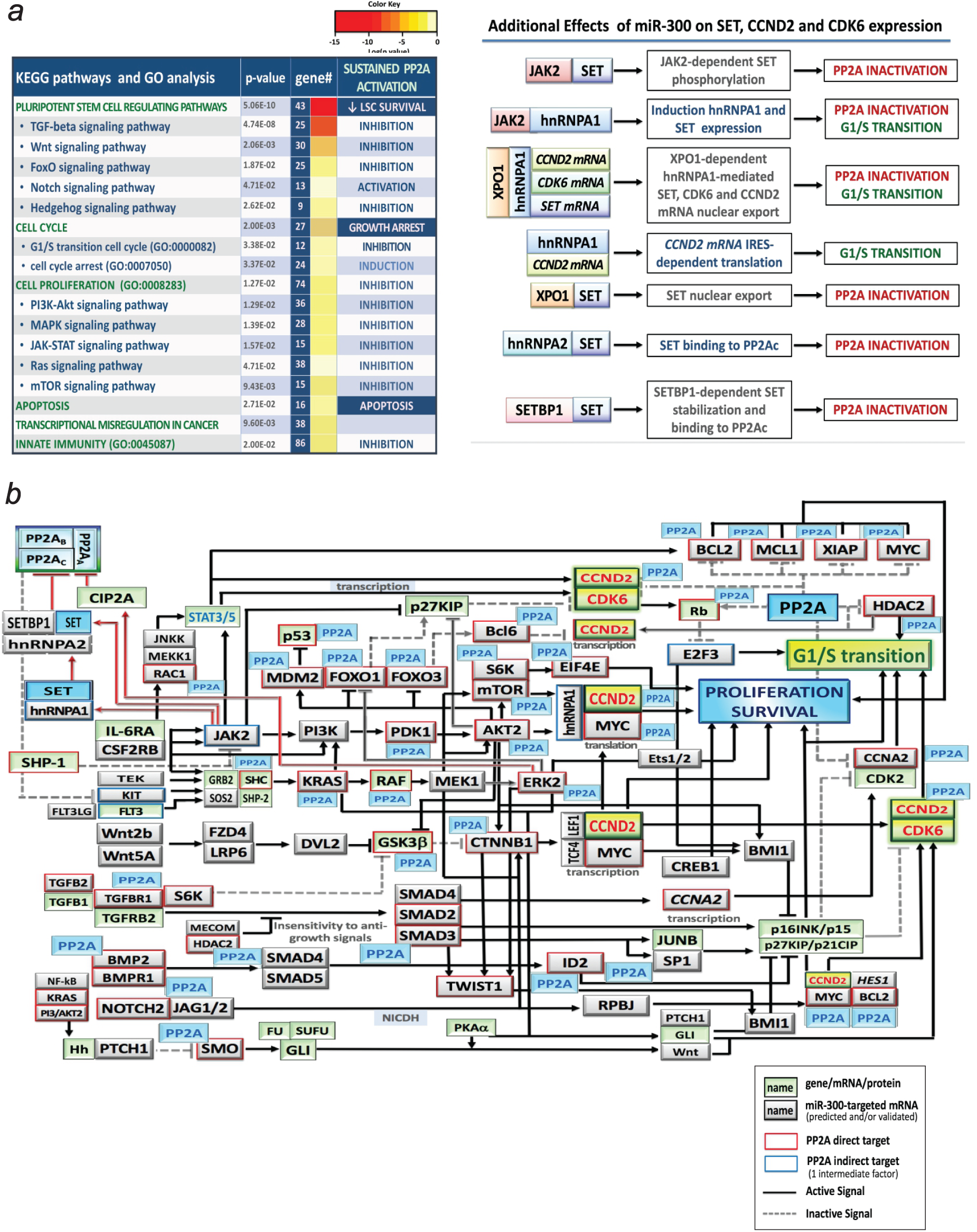
MIR300 is a master PP2A activator and an inhibitor of G1/S transition. (**a**) (*left*) KEGG/GO analysis of *MIR300* effects on signal transduction pathways; (right) additional levels at which *MIR300* exerts its inhibitory effects on SET, CCND2 and CDK6. (**b**) Pleiotropic role of PP2A on the activity of the predicted *MIR300*-targeted factors.

**Supplementary Figure 3.**
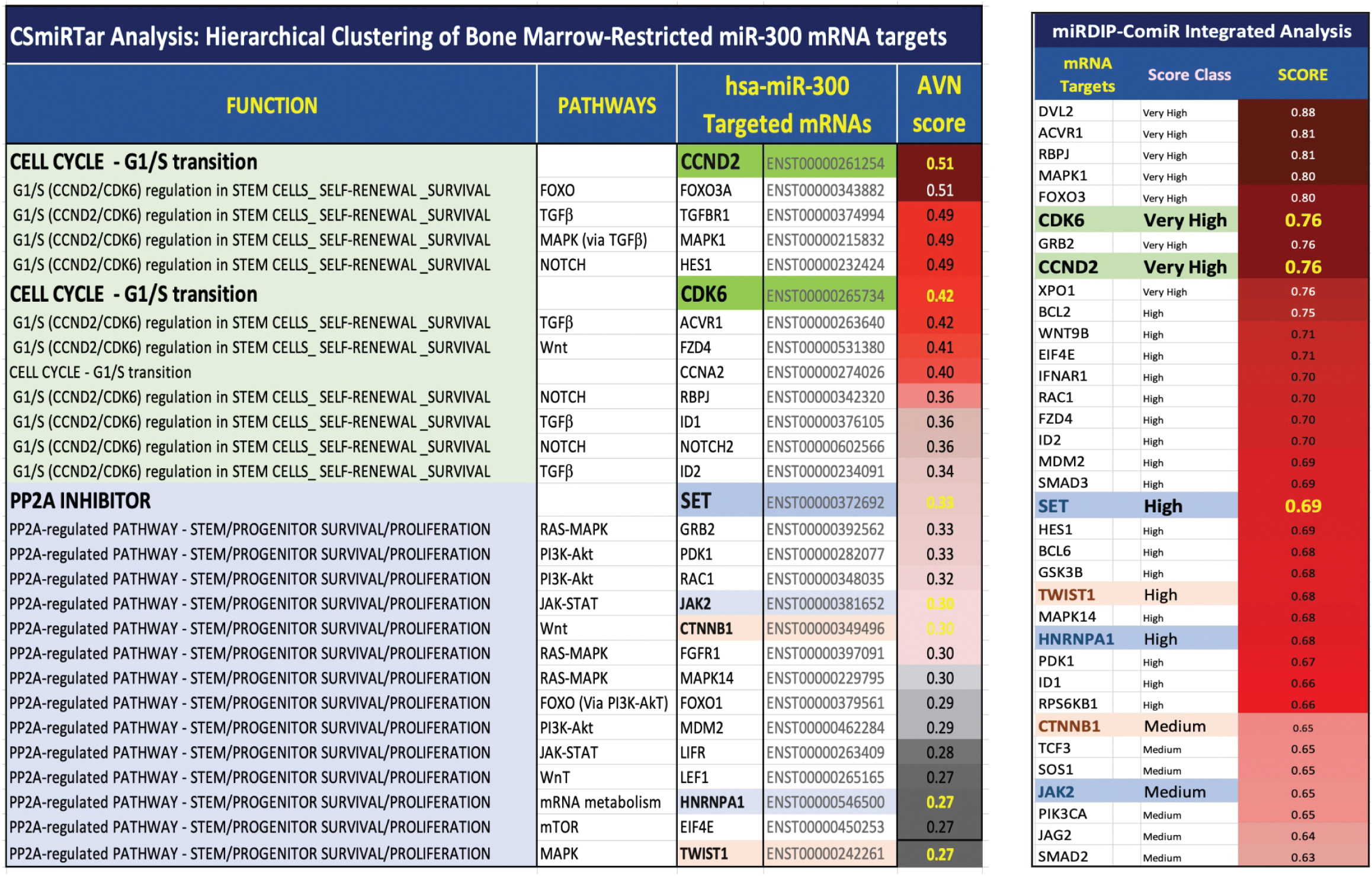
MIR300 dose-dependent target selection. CSmiRTar (filters: bone marrow normal and myeloid leukemia cells) and miRDIP-ComiR integrated analyses show functional clustering of predicted/validated *MIR300* targets adjusted according to *MIR300* levels required for their inhibition and expression of *MIR300* targets in bone marrow and leukemic cells.

**Supplementary Figure 4.**
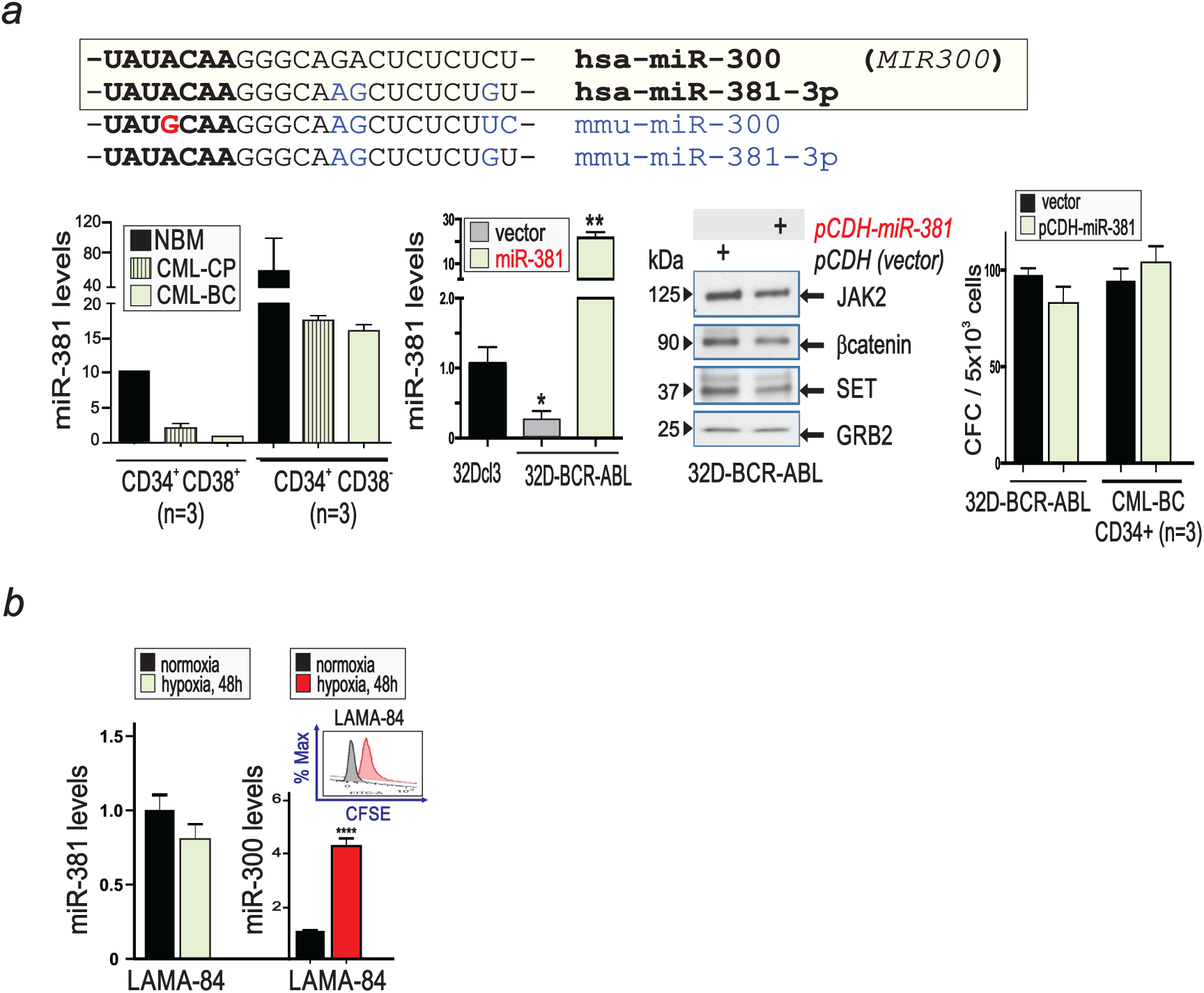
miR-381-3p expression and activity in leukemic stem and progenitor cells. (**b**) (*top*) Sequence homology between human hsa-*MIR300*-3p (*MIR300*) and intragenic hsa-miR-381-3p and between mouse mmu-*MIR300*-3p (*MIR300*) and mmu-miR-381-3p mature microRNAs; seed sequences are in bold and mismatching bases in blue. The mismatch at +4 position between human and mouse *MIR300* is depicted in red. (*bottom left to right*) miR-381-3p levels in CML and NBM CD34^+^CD38^-^ and progenitor CD34^+^CD38^+^ cells, 32Dcl3, vector and miR-381-3p-transduced 32D-BCR-ABL1 myeloid precursors; effect of miR-381-3p on *MIR300* targets and on clonogenic potential (CFC) of vector and miR-381-3p-expressing 32D-BCR-ABL and CD34^+^ CML-BC (n=3) cells. (∗*P*<0.05, ∗∗*P*<0.01). (**c**) miR-381-3p and *MIR300* levels in LAMA-84 cells cultured in normoxic and hypoxic conditions; *inset:* effect of hypoxia on CFSE^+^LAMA-84 cell division.

**Supplementary Figure 5.**
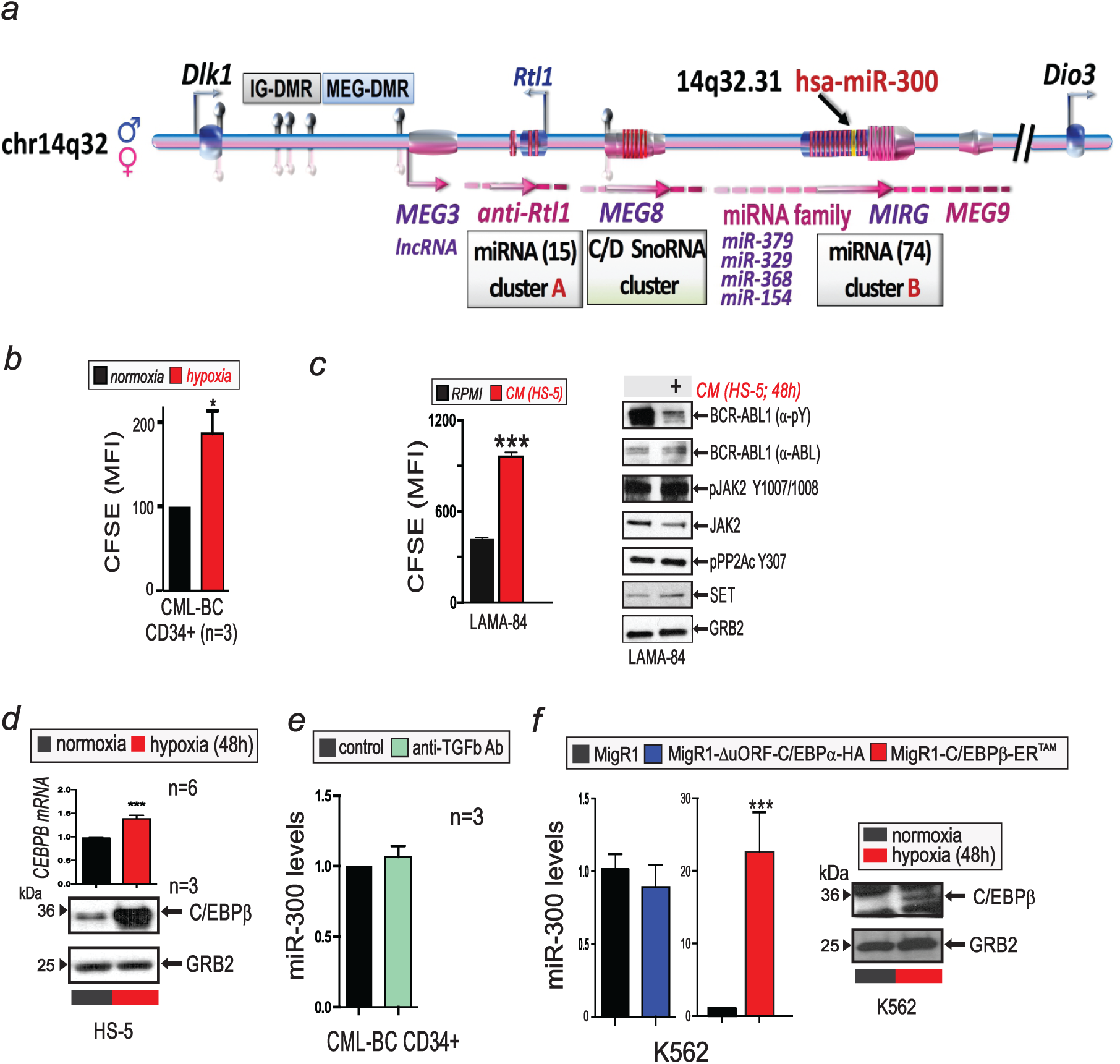
BMM-induced C/EBPβ-dependent MIR300 tumor suppressor activity preserves LSCs. (**a**) Structure of 14q32 DLK1-DIO3 genomic imprinted locus hosting the MEG3-regulated hsa-*MIR300*. (**b**) Effect of hypoxia on proliferation of CFSE^+^CD34^+^ CML-BC cells. (**c**) Effect of MSC (HS-5)-derived CM on LAMA-84 proliferation expressed as fold changes of CFSE mean of fluorescence intensity (MFI)±SEM and on: BCR-ABL1 expression (*α*-ABL1) and activity (*α*-PY), phospho-BCR-ABL1, JAK2 expression and activity JAK2 Y1007/1008, PP2A activity (pPP2AY307 inactive form) and GRB2 used as a control (blots are representative of three independent experiments). (**d**) Levels of C/EBP*β* and GRB2 mRNA and protein in HS-5 cells exposed to hypoxia (48h; 1% O2). (**e**) Effect of neutralizing TGF*β* antibody (anti-TGF*β* Ab; 48h, 1.25 μg/ml) on *MIR300* levels in CD34^+^ CML-BC cells. (**f**) Effect of ectopic C/EBP*α* (MigR1-*Δ*uORF-C/EBP*α*-HA) and C/EBP*β* (MigR1-C/EBP*β*-ER^TAM^) on *MIR300* levels in K562 cells. Immunoblot shows levels of C/EBP*β* and GRB2 in normoxic and hypoxic K562 cells.

**Supplementary Figure 6.**
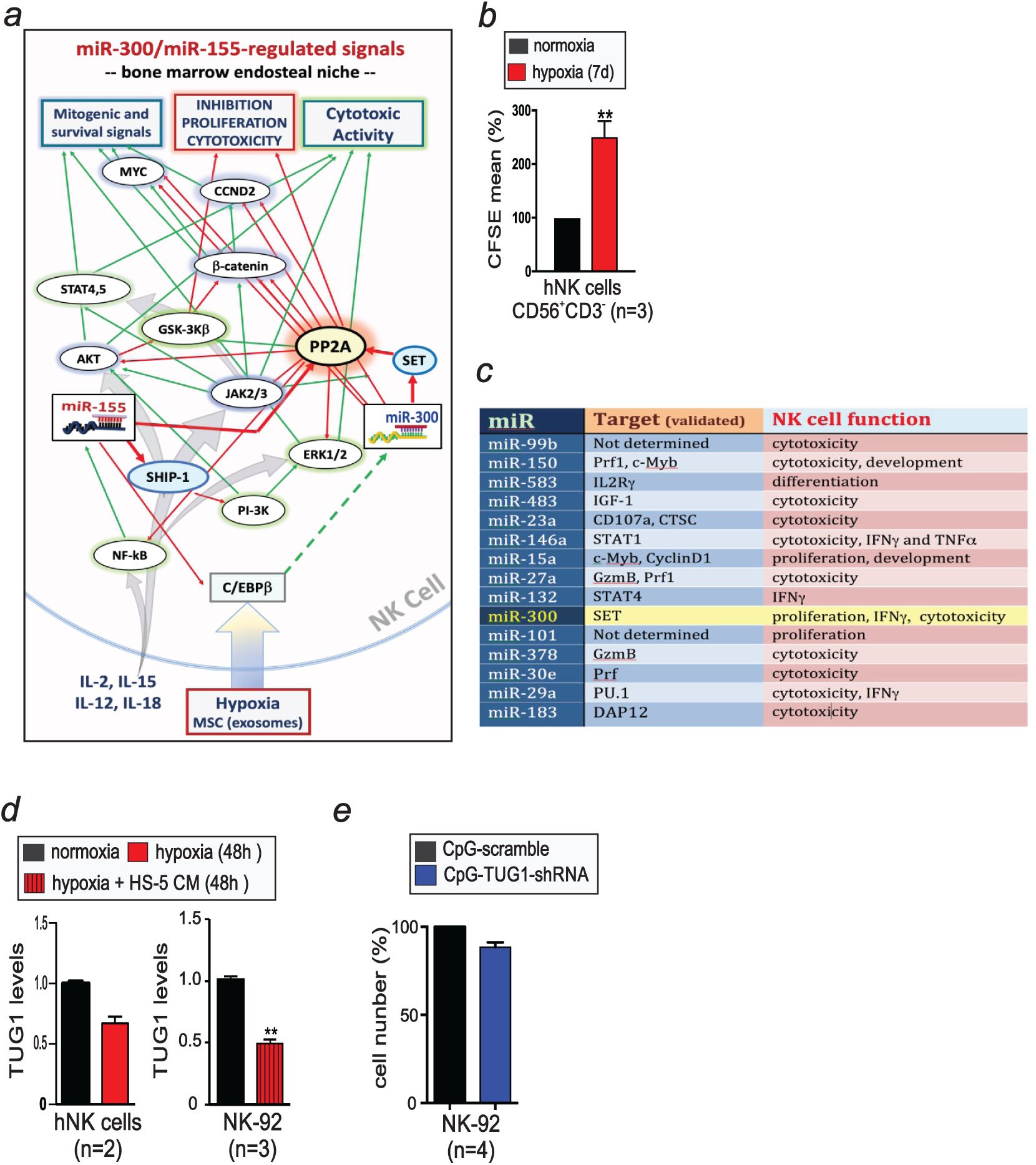
MIR300 inhibits NK cell growth and activities. (**a**) Validated *MIR300* and miR-155 pathways predicted to occur in the bone marrow endosteal niche. (**b**) Effect of hypoxia (1% O_2_, 7 days) on CFSE^+^NK cell proliferation (% CFSE mean of fluorescence). (**c**) HS-5 exosomal miRNAs described as negative regulators of NK cell proliferation/activity and their experimentally validated targets. miRNA RNAseq was performed on an Illumina platform using libraries derived from 100 ng RNA/sample from HS-5 exosome purifications (n=3). (**d**) Effect of hypoxia (1% O2) and HS-5 CM (48h) on TUG1 expression in CD56^+^CD3^-^ primary NK and NK-92 cells. (**e**) Effect of *CpG-TUG1-shRNA* and *CpG-scramble* (200-500 nM, 5 days) on IL-2-induced NK cell proliferation (% cell number). Data are represented as mean±SEM.

**Supplementary Figure 7.**
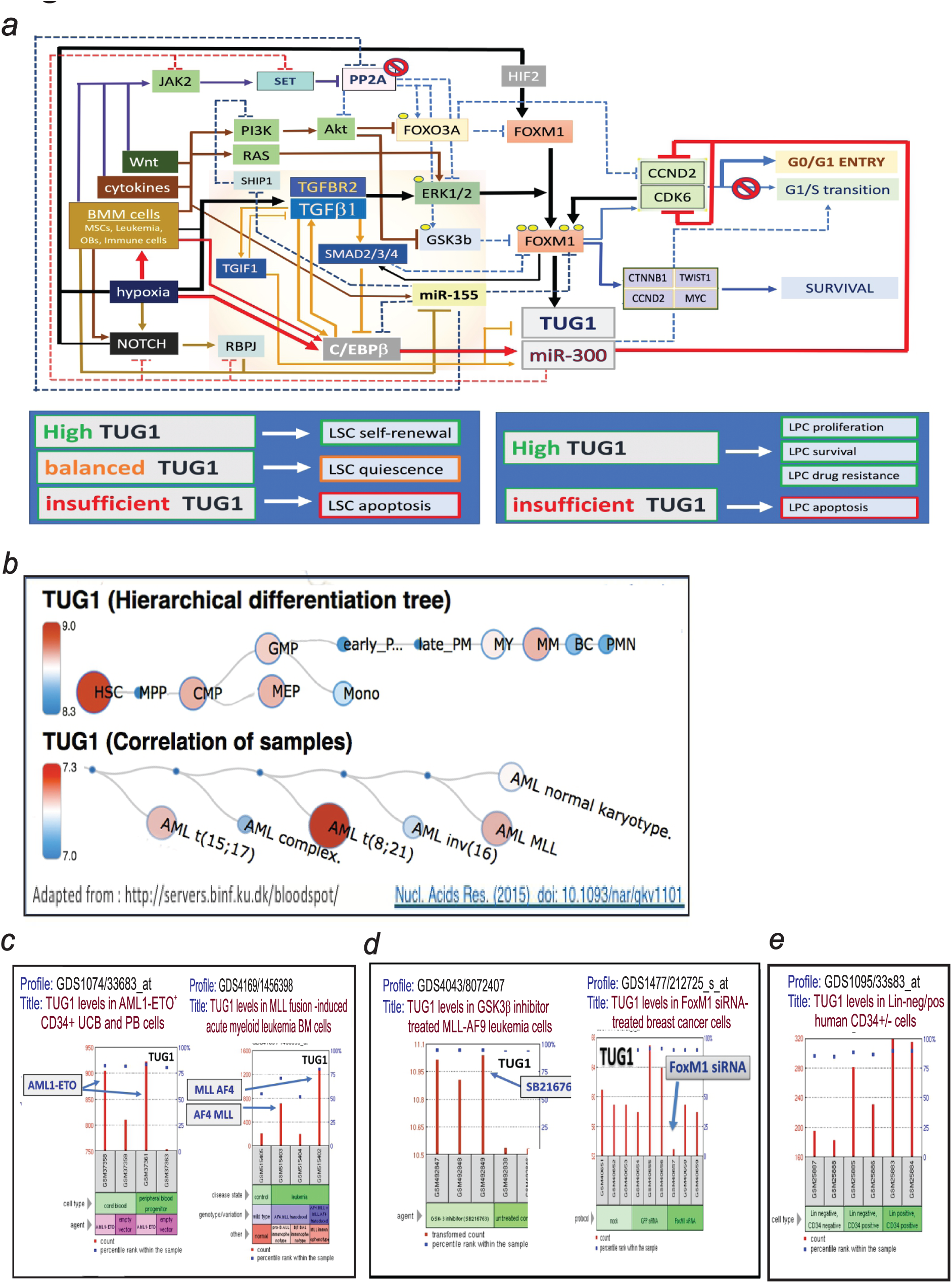
TUG1 selectively allows MIR300 anti-proliferative activity in LSCs. **(a)** Graphic representation of the predicted signaling network controlling LSC quiescence and survival through the BMM-C/EBP*β*-*MIR300* and BMM-TGF*β*-FoxM1 pathways. Dotted lines indicate inactive pathways, line thickness indicates relevance of the signal for LSC quiescence. Red lines indicate signals that are going to increase *MIR300* levels. Black lines signals that are going to increase TUG1 levels. (bottom) effects of different TUG1 levels on leukemic stem (LSC) and progenitor (LPC) cell fate. **(b)** BloodSpot array-based TUG1 expression levels during normal myelopoiesis and in different AML subtypes. GEO Profiles show TUG1 levels **(c)** in AML1-ETO- and MLL-AF4-expressing human CD34^+^ UCB/PB and mouse AML cells, and **(d)** (*left*) in GSK3*β* inhibitor-treated MLL-AF9 AML cells, and (*right*) upon siRNA-mediated FoxM1 downregulation in breast cancer cells. **(e)** TUG1 levels in lineage-negative (Lin^-^) and - positive (Lin^+^) CD34^-^ and CD34human stem/progenitor cells from healthy individuals.

**Supplementary Figure 8.**
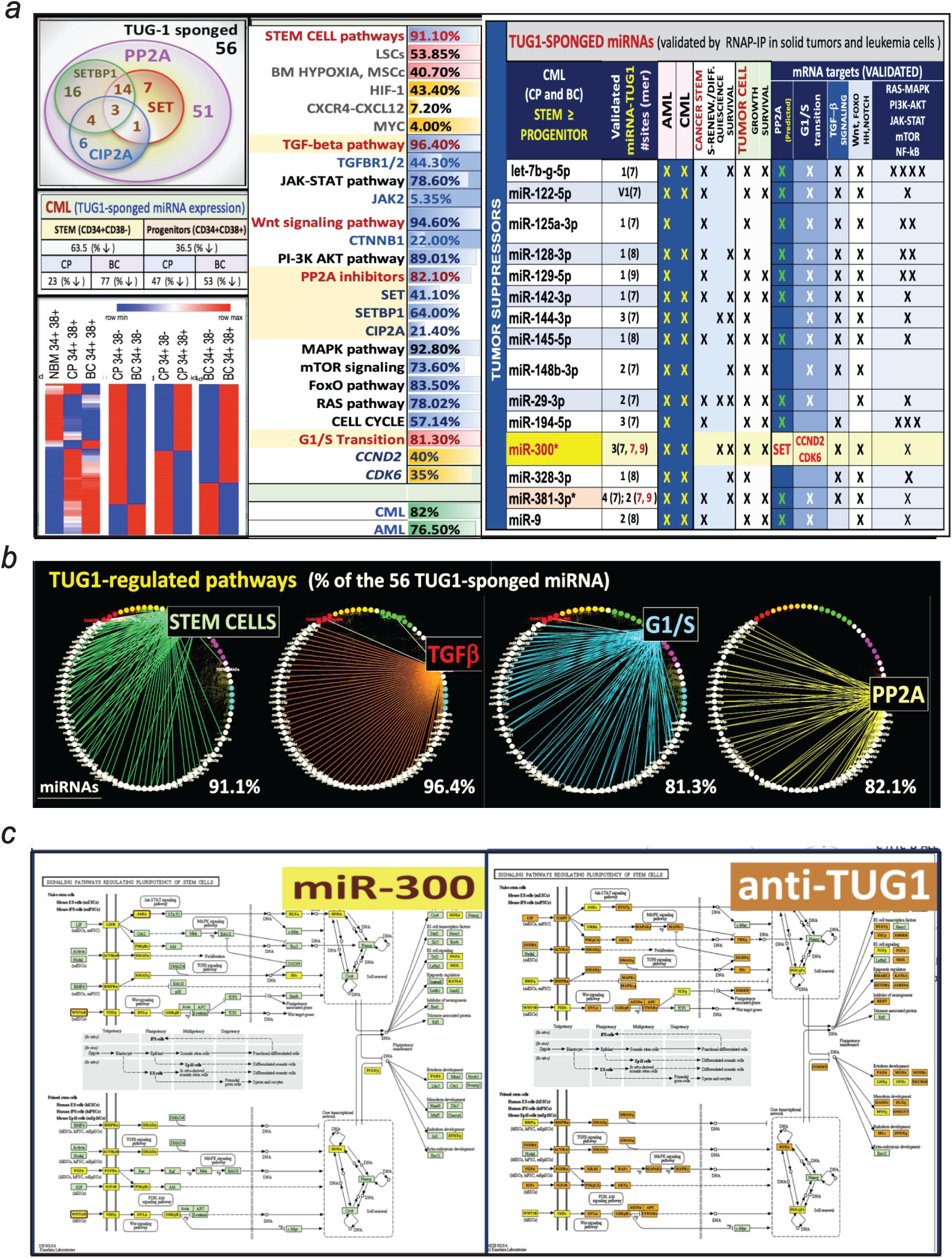
Analysis of TUG1-sponged miRNAs and their mRNA targets. **(a)** (*left-to-right*): Integration of predicted TUG1-sponged miRNAs with RNAseq data (heatmaps) from BM CD34^+^CD38^-^ and CD34^+^CD38^+^ CML (CP and BC) and NMB (n=3/group). Effect of predicted TUG1-sponged miRNA on PP2A inhibitors (Venn diagram); Functional integration of TUG1-sponged miRNAs and their mRNA targets into regulatory network using DIANA Tools (http://diana.imis.athena-innovation.gr), and assessment of their biological function by KEGG/GO analysis. **(b)** Circular diagrams show top 4 pathways affected by TUG1-sponged miRNAs. **(c)** Predicted effect of *MIR300* and of TUG1-sponged miRNAs (anti-TUG1) on Pluripotent Stem Cell Signaling pathways. <u>Method:</u> miRNA RNAseq was performed with 100 ng RNA/samples (HS-5 exosomal RNA purifications, n=3; and, LSC-enriched CD34^+^CD38^-^ and progenitor CD34^+^CD38^+^ cell fractions from BM frozen specimens of CML-CP and -BC, n=3 patients and healthy individuals (NBM) per group) and indexed small RNA libraries were prepared using a modified version of the TruSeq Small RNA Library preparation kit (Illumina) and final library QC was performed using the LabChip XT (Caliper Life Sciences). The libraries were pooled and sequenced on an Illumina MiSeq 50bp run. Following standard data processing and demultiplexing, adaptor trimming was performed using Trimmomatic.

